# Nodule microbiome functions shape the performance of a wild perennial legume

**DOI:** 10.64898/2026.04.06.716301

**Authors:** Moshe Alon, Guy Dovrat, Yoni Waitz, Amir Erez, Efrat Sheffer, Omri Finkel

## Abstract

Nitrogen fixing legume nodules are typically viewed as the product of a bilateral mutualism between host plants and nitrogen-fixing rhizobia, yet nodules also harbor diverse non-rhizobial endophytes whose functional significance remains poorly understood, especially in wild legumes and uncultivated soil. Here, using the wild Mediterranean shrub *Calicotome villosa*, we performed a soil inoculation experiment to test whether plant performance is linked to the functional composition of the nodule microbiome. Soil inocula from different natural sites produced strong differences in nodulation success, plant biomass, leaf nitrogen concentration, nitrogen fixation rate, and nodule allocation under otherwise uniform conditions. Although *Bradyrhizobium* dominated all nodules, species composition varied among inoculation sources, and non-rhizobial endophytes reached substantial abundance in some treatments. Functional profiles of the nodule microbiome were significantly associated with plant phenotype, with the strongest coupling observed for traits related to nodule investment. Targeted and genome-wide analyses identified trait-associated genes in both symbionts and endophytes, including genes involved in nitrogen cycling, ammonium transport, denitrification, pyrimidine degradation, sulfur assimilation, and type VI secretion systems. Several of these functions were not part of the canonical symbiosis machinery, yet were strongly associated with plant nitrogen status, biomass accumulation, or nodule mass fraction. Together, our results show that legume performance is better predicted by the collective functional composition of the nodule microbiome than by the primary symbiont alone. These findings support a broader view of nodules as multipartite microbial communities.

## Introduction

Symbiotic dinitrogen (N-_2_) fixation in terrestrial plants arises from a mutualistic interaction between plants, mainly legumes (Fabaceae), and N-2-fixing bacteria, mainly rhizobia. This mutualism, occurring in specialized root nodules, supplies most of the naturally bioavailable nitrogen in the terrestrial biosphere [1, 2]. Rhizobial symbionts belong predominantly to the Alphaproteobacteria, with *Bradyrhizobium*, *Rhizobium*, and *Ensifer* being the most prominent genera in terms of isolation frequency, host range, and agricultural importance [3–8].

Some rhizobia are less efficient N_2_ fixers than others, and some do not fix nitrogen at all [9–12]. The stability of the legume-rhizobia symbiosis is maintained through host control mechanisms that limit exploitation by less mutualistic strains. These mechanisms operate both during partner choice, when the host screens for potential symbionts, and after infection, when the plant sanctions non-performing nodules by restricting resources like oxygen and carbon [13–16]. Yet given the diversity of bacteria that can colonize nodules, including strains that fix little or no nitrogen, understanding the full range of symbiotic outcomes requires looking beyond the dominant rhizobial symbiont.

Although individual infection threads and the resulting nodules are often dominated by a single rhizobial strain, multiple bacterial cells, and sometimes multiple strains, can contribute to nodule formation, and nodules commonly host diverse non-rhizobial endophytes (NREs) [17–19]. These NREs are consistently detected at low abundances alongside rhizobia [20, 21]. Within nodules, NREs can influence host performance in multiple ways, affecting plant biomass and nitrogen content while also improving stress responses [19]. Most studies to date found that NREs tend to improve symbiotic performance, including nitrogen fixation efficiency, plant biomass, and plant nitrogen content [22–24]. But detrimental effects, including reduced shoot biomass and plant fitness, have also been observed [25, 26].

Taxonomically, NRE communities are often dominated by genera such as *Bacillus*, *Pseudomonas*, *Paenibacillus*, *Enterobacter*, and *Agrobacterium,* and they harbor genes associated with a wide range of plant growth-related functions, including indole-3-acetic acid production, phosphate solubilization, siderophore production, 1-aminocyclopropane-1-carboxylate (ACC) deaminase activity, and even active nitrogenase genes [19, 27–29]. NREs can also mitigate abiotic stresses such as salinity, heat, drought and metal contamination [23, 30, 31]. Yet studies that examined NREs have focused almost exclusively on agricultural legumes such as soybean, chickpea and pea, and the identity and functional potential of NREs in wild legumes under natural conditions remains unknown.

To date, most nodule microbiome research has relied on culture-dependent assays or on 16S rRNA amplicon sequencing [19, 32, 33]. While culture-based methods allow direct testing of traits such as phosphate solubilization, they miss the unculturable majority of microbes. Conversely, amplicon sequencing provides a taxonomic view of bacterial community composition but little functional information. Shotgun metagenomics, which can capture the functional potential of the entire microbial community, remains rare in studies of the N-_2_-fixing symbiosis, largely because of the technical challenges posed by host DNA contamination (but see [34, 35]). Here, we characterize the structure and functional repertoire of the legume nodule microbiome and provide a detailed, culture-independent view of this complex multipartite symbiosis.

We used *Calicotome villosa*, a widespread shrub from the Mediterranean basin, that occurs across diverse soil types, altitudes and precipitation regimes, to study how nodule microbiome composition and function influence plant performance. *Calicotome villosa* is associated with a single rhizobial genus, *Bradyrhizobium* [36]. It shows a strong seasonal pattern of nitrogen fixation [37], and tightly regulates allocation to nodule biomass, measured as nodule mass fraction (NMF; nodule mass divided by total plant mass), in response to soil water and nitrate availability [38, 39]. We therefore asked whether variation in nodule microbiome composition and function predicts plant performance, which bacterial genes and pathways are associated with whole-plant and nodule-related traits, and whether these associations differ between the primary symbiont and NREs.

To test the relationship between nodule microbiomes and plant performance, we grew *C. villosa* seedlings with inocula derived from a diverse set of natural soils. We hypothesized that plant performance would vary as a function of both the colonizing symbiont and its co-occurring NREs. To characterize nodule microbial taxa, and identify microbial genes and pathways associated with plant performance, we integrated metagenome-assembled genomes (MAGs), read-based KEGG Orthology (KO) annotations, targeted analyses of nitrogen metabolism genes, and machine-learning analyses. By classifying genes and MAGs as belonging either to the primary rhizobial symbiont (*Bradyrhizobium*) or to NREs, we assessed their relative associations with plant traits. We found that plant performance was strongly linked to nodule functional profiles derived from both the symbiont and its accompanying NREs. We show that microbial genes not directly involved in canonical symbiosis can nonetheless have substantial effects at the whole-plant scale.

## Methods

### Controlled experiment

We conducted a controlled pot experiment to evaluate how soil-derived microbial inocula influence nodulation and growth in *Calicotome villosa* seedlings. Seeds were collected from natural populations at three field sites (Ramat-Hanadiv, Carmel, and Caesarea; Table S1), pooled across populations to minimize genotypic effects, surface-sterilized, and germinated in agar plates in an incubator (12-12 light-dark, 20 °C) using protocols developed to synchronize germination. Germinated seedlings were transplanted to 200 ml pots filled with sterile, inert perlite (Agrekal, 0.2 mm, density 2.2 g cm^-3^).

All plants grew under uniform environmental conditions in a net house with a transparent rain-shed roof and net walls at Beit Dagan, Volcani Center, during the natural growing season (autumn to spring). Seedlings were initially fertigated with a complete nutrient solution (N:P:K = 70:20:60 with all necessary micronutrients). After 38 days, all pots were washed thoroughly with water to remove residual fertilizer, and plants were divided into ten treatments (*n* = 10 pots per treatment). Nine treatments were inoculated by spreading 6 ml of sieved soil, collected from one of nine different locations, on the perlite surface (Table S1, Fig. S1). Seven inoculation soils were collected from beneath adult *C. villosa* canopies at different field sites; one was from beneath *Genista fasselata*, a co-occurring N2-fixing shrub (Carmel site); and one was from Yatir, a semi-arid site outside the distribution of *C. villosa*. A tenth treatment served as an uninoculated control. All soils were collected within 48 hours of application, in early winter, and stored at room temperature until use. For the remaining four months, all seedlings received a nitrogen-free nutrient solution (N:P:K = 0:20:60) with all necessary micronutrients, including molybdenum, to ensure that nitrogen was available only through biological fixation.

### Measurements of plant performance traits

At harvest (day 127), all surviving plants (*n* = 8-10 per treatment; *N* = 85) were measured for height, above- and belowground dry biomass, root nodule biomass, and nitrogen fixation rate. Fixation was estimated by acetylene reduction assay (ARA): root segments with attached nodules were incubated with 10% acetylene, and ethylene production was quantified by GC-FID (Focus, Thermo-Finnigan) from gas samples taken at 30 and 60 minutes. Fixation rate was expressed per gram of incubated nodule biomass. Nodule vitality was assessed by internal color (pink-red). All biomass was oven-dried at 60 °C for 72 hours. Root mass fraction (RMF) and nodule mass fraction (NMF) were calculated from these measurements. Leaf carbon and nitrogen concentrations were measured using an elemental analyzer (FLASH2000, Thermo Scientific).

### Analyses of inoculation soils

Inorganic nitrogen (NO₃⁻-N and NH_₄⁺_-N) was extracted with 2M KCl and measured spectrophotometrically. Organic carbon and total nitrogen were measured by elemental analysis after carbonate removal. Nitrogen accumulated in plant biomass over the experiment exceeded the nitrogen content of the soil inoculum by 375-1,103-fold, confirming that inoculation-derived nitrogen was negligible (Table S1).

### Nodule sampling and DNA extraction

Multiple nodules per plant were collected from each treatment and surface-sterilized by sequential immersion in 70% ethanol (30 s), 3% sodium hypochlorite (2 min), and five sterile water washes. Sterilized nodules were crushed by bead-beating and extracted for genomic DNA using the PowerSoil kit (Qiagen). Shotgun metagenomic sequencing was performed on the Illumina NovaSeq X Plus platform (Novogene) to a depth of 25-46 million 150 bp paired-end reads per sample (mean ∼34 million). Sequence data are available at NCBI BioProject PRJNA1438069.

### Metagenomic analysis

Reads were quality-filtered using Trimmomatic and validated with BBTools [40, 41]. We combined assembly-free and assembly-based approaches. In the assembly-free approach, taxonomic profiles were generated using SingleM with the GTDB r226 database, and reads were functionally annotated by DIAMOND BLASTX against the KEGG Prokaryotes database (e-value ≤ 1e-5)[42]. Raw KO counts were normalized by estimated genome number using 106 bacterial single-copy genes [43]. In the assembly-based approach, reads were assembled with MetaSPAdes and binned using five algorithms (MetaBAT2, MaxBin2, CONCOCT, COMEBin, SemiBin2), integrated with Binette [44–49]. Bin quality was assessed with CheckM2, and only MIMAG-compliant MAGs (≥50% completeness, ≤10% contamination) were retained [50, 51]. MAG taxonomy was assigned using GTDB-Tk (release 226), and relative abundances were quantified using CoverM. Eukaryotic contamination was assessed using EukFinder. Full pipeline details, including MAG annotation with Prodigal and MicrobeAnnotator, phylogenetic reconstruction in Anvi’o, and co-occurrence analysis with FastSpar, are provided in Supplementary Methods [52–57]. The metagenomic pipeline is available at github.com/Naturalist1986/Metagenomic_Assembly_and_Binning_pipeline.

### Statistical analysis

We used Procrustes analysis and Mantel tests (vegan R package) to evaluate congruence between plant trait ordinations and nodule KO functional composition [58–60]. To identify bacterial genes associated with plant performance, we performed (i) a targeted analysis of 87 nitrogen metabolism KOs using univariate linear regression with multi-method FDR correction, and (ii) an unrestricted Boruta feature selection across all KOs [61]. For each significant KO, taxonomic origin was determined from the MAG data to distinguish symbiont- from NRE-associated genes. Full details of ordination methods, transformation approaches, filtering criteria, and correction strategies are provided in Supplementary Methods. Analysis scripts are available at github.com/Naturalist1986/nodule-microbiome-analysis-scripts.

### Global Bradyrhizobium biogeography

To contextualize the *Bradyrhizobium* clades identified in our experiment, we analyzed their abundance across 1,963 global soil metagenomes in relation to 18 soil and climatic variables (Supplementary Methods).

## Results

### The phenotypic response of *C. villosa* to inoculation is site-specific and tracks nodule microbial functions

To test how soil-derived microbial inocula affect legume nodulation and growth, we established a controlled experiment in which the perennial legume *C. villosa* was grown in fertilized perlite under uniform conditions. After 38 days of growth, we inoculated the perlite surface with 3% (v/v) sieved soil collected from eight different sites (Fig. S1, Table S1). Nodulation varied markedly among inoculation treatments. Among plants inoculated with soil collected from beneath *C. villosa* shrubs, 80-100% formed nodules. By contrast, soils from sites where *C. villosa* does not occur naturally (YTIR and DGN) did not induce root nodules, whereas soil collected from beneath a different plant species, *G. fasselata* (CAR-G), induced nodulation in 40% of individuals (Fig. 1A). These results point to a plant-soil feedback and support a link between host and symbiont biogeography: soil collected from beneath the host was the strongest inoculant.

**Figure 1:**
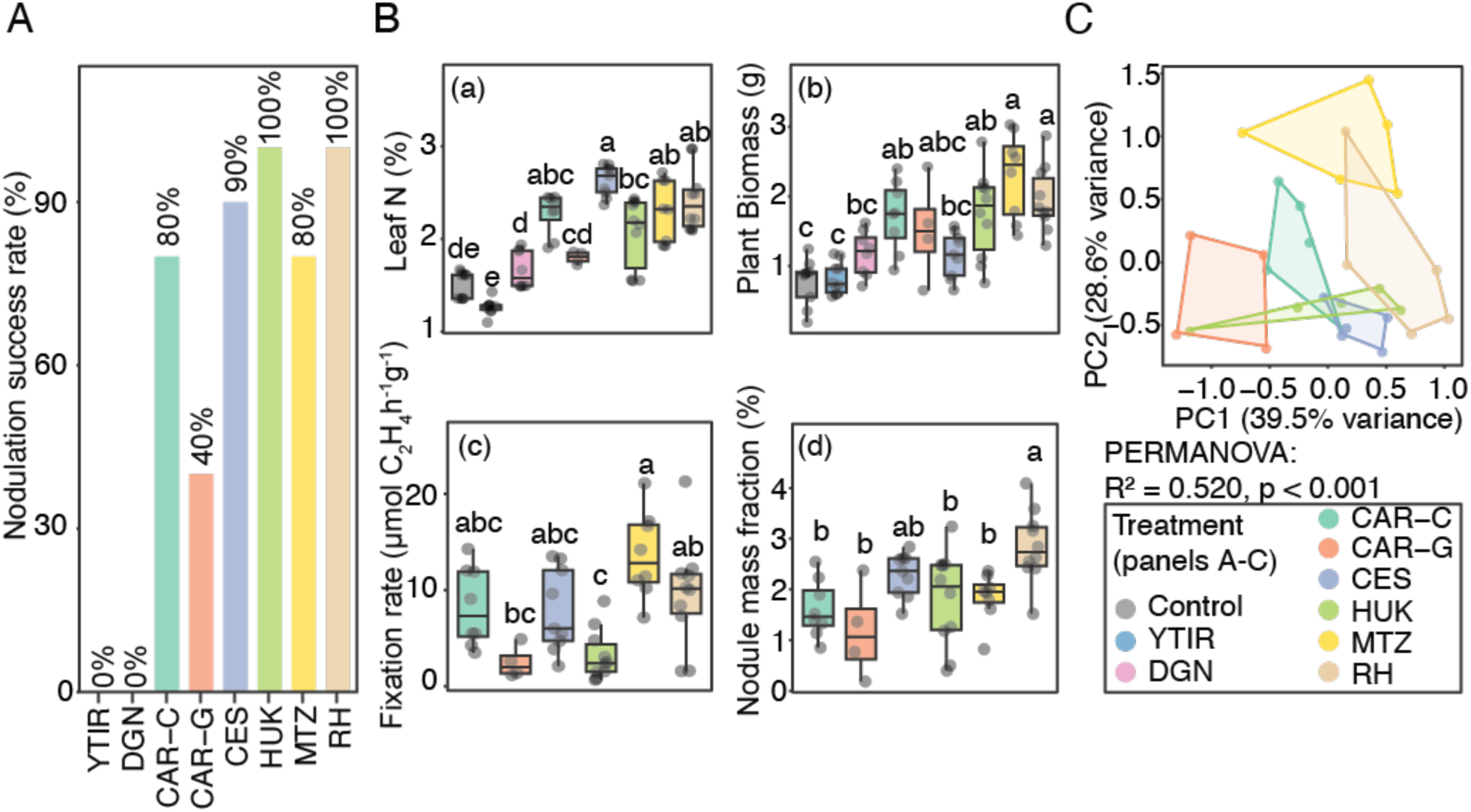
Influence of inoculation source on plant performance traits. **(A)** Nodulation success (% of plants nodulated) of *C. villosa* seedlings inoculated with soil from the different sites. **(B)** Plant properties under the different soil inoculations. Boxplots display (a) leaf nitrogen concentrations (%), (b) total plant biomass (g), (c) nitrogenase activity (fixation rate per nodule dry mass; μmol C_2_H_4_ h^-1^g^-1^), and (d) nodule mass fraction (NMF; %). Horizontal lines within boxes represent the median; box limits indicate the 25th and 75th percentiles; whiskers extend to 1.5 times the interquartile range; and individual data points are overlaid. Statistical significance was determined by ANOVA followed by Tukey’s HSD post-hoc test (p < 0.05), shown using the compact letter display. **(C)** Principle component analysis (PCA) on the normalized four plant traits. Polygons define the convex hull encompassing all samples within each inoculation treatment. Statistical significance was determined by a PERMANOVA test.

As expected, nodulation increased both leaf nitrogen concentration and plant biomass, although the magnitude of this effect varied among inoculation treatments (Fig. 1B). Among nodulating plants we also observed significant variation in fixation rates and in nodule mass fraction. More broadly, plant traits clustered by soil inoculation, with the first two ordination axes together explaining 68.1% of variance (Fig. 1C). Together, these results show that soil-derived nodulation and microbiota drive distinct physiological states in *C. villosa* under otherwise uniform abiotic conditions.

### Nodule microbiome composition is highly site-specific

Given the differences in plant performance across soil inoculation treatments, we next asked how nodule microbiome composition varied among them. We performed shotgun metagenomic sequencing on crushed, surface-sterilized nodules from all treatments and annotated individual sequence reads for both taxonomy and function. As expected, *Bradyrhizobium*, the primary symbiont genus of *C. villosa*, dominated all samples with 16 different species identified within the genus (Fig. 2A). *Bradyrhizobium* accounted for 91-95% of reads in most samples, except in the MTZ treatment, where it comprised only 80% of the nodule microbiome. Non-rhizobial endophytes (NREs) were taxonomically diverse, particularly in the MTZ treatment, with Burkholderiales reaching the highest relative abundance among them (Fig. 2A).

**Figure 2:**
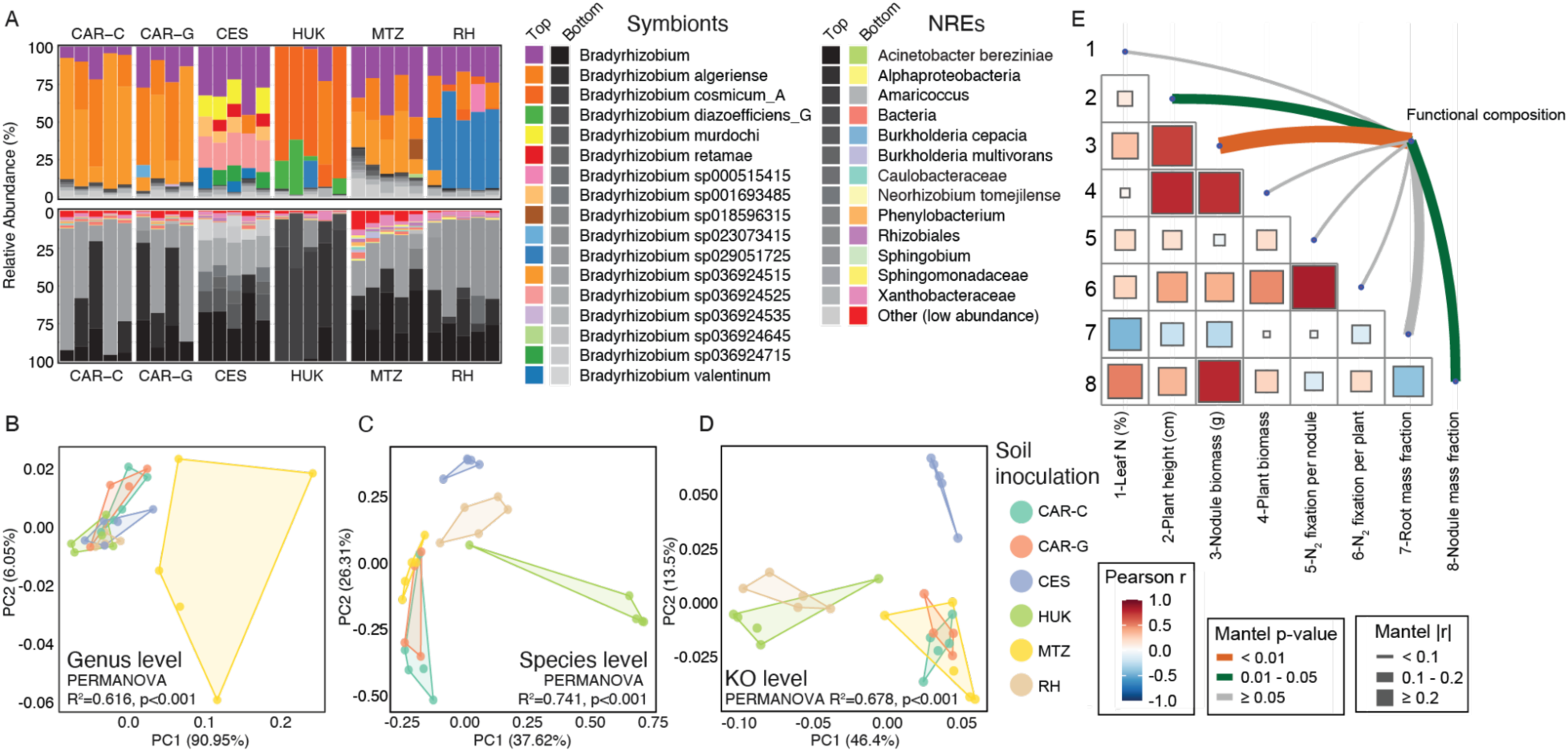
Assembly-free taxonomic and functional composition of the nodule microbiomes. **(A)** Relative abundance barplot of top 30 assigned taxonomic groups at multiple levels; The taxa displayed are the lowest taxonomic level assigned by SingleM. Both barplots show the same data with the top plot showing NREs in grayscale and *Bradyrhizobium* in color, and the bottom is inverted - NREs in color, and *Bradyrhizobium* in grayscale. **(B-C)** PCoAs based on genus **(B)** and species **(C)** composition based on SingleM derived taxonomy. Polygons define the convex hull encompassing all samples within each inoculation treatment. **(D)** PCoA of KO functional composition (Hellinger + Bray-Curtis distances). Statistical significance was determined by a PERMANOVA test. **(E)** Mantel test of KEGG ortholog (KO) composition in nodules vs. plant properties. The lower triangle of the figure displays a Pearson correlation heatmap among all eight measured plant traits. Curved lines represent the Mantel test results comparing dissimilarity in each trait to dissimilarity in microbial function composition: line colors encode significance category (p < 0.01, 0.01 ≤ p < 0.05, or p ≥ 0.05), line width encodes the magnitude of the Mantel statistic (|r| < 0.1, 0.1–0.2, or ≥ 0.2).

Soil source strongly influenced both taxonomic and functional composition of the nodule microbiome. PCoA of taxonomic composition showed that, at the genus level, only nodules from the MTZ treatment were clearly separated from the other treatments, consistent with their relatively high abundance of NREs (Fig. 2B). At the species level, however, nearly all sites were clearly separated (Fig. 2C), indicating that each site supported a distinct combination of *Bradyrhizobium* species. Soil source also explained much of the variance in functional composition, with the first two axes accounting for 59.9% of the variance (Fig. 2D).

To test whether microbiome functional composition was associated with plant phenotype, we compared the ordination of microbial functional profiles (Fig. 2D) with the ordination of plant traits (Fig. 1C) using Procrustes analysis. This revealed a significant correlation between microbiome function and plant trait profiles (r = 0.52, m² = 0.73, p < 0.001; Fig. S2A). We further evaluated this relationship by using Mantel tests between Euclidean distances calculated from eight plant traits and Bray-Curtis dissimilarities derived from metagenomic KO profiles (Fig. 2E). All associations were positive, indicating that greater phenotypic dissimilarity among plants corresponded to greater dissimilarity in microbiome functional composition. Nodule biomass exhibited the strongest and most significant association (Mantel r = 0.316, p = 0.002), suggesting that investment in nodules is the plant trait most tightly coupled to nodule microbiome. Notably, nodule biomass was also strongly correlated with plant height, plant biomass and consequently NMF. Plant height (Mantel r = 0.152, p = 0.024) and NMF (Mantel r = 0.14, p = 0.029) also showed significant associations with microbiome functional composition, consistent with a broader link between symbiotic investment and microbiome function. Together, these results indicate that plant phenotype, and especially investment in nodules, is closely associated with the functional profile of the nodule microbiome.

### *C. villosa* nodule *Bradyrhizobia* symbionts in the context of global *Bradyrhizobium* biogeography

To resolve the genomes of nodule residents, we assembled and binned the metagenomic data. This yielded 44 MAGs, 15 of which belonged to the primary symbiont, *Bradyrhizobium*. We then mapped reads back to the MAGs to estimate their abundance within each community. This assembly-based approach also helped separate bacterial signals from host contamination. Although 92-97% of reads mapped back to the de novo assemblies, only a subset mapped to bacterial contigs (45.77% in MTZ to 88.84% in HOK), indicating substantial host or fungal contamination in some samples (Fig. 3A). Across all soil inoculation treatments, nodules harbored at least two distinct *Bradyrhizobium* MAGs per treatment, along with lower-abundance MAGs from non-rhizobial endophytes, including *Cellvibrio*, *Novosphingobium*, *Burkholderia*, and members of the Caulobacteraceae, which we collectively classify as NREs (Fig. 3B-C, Fig. S2B). These NRE MAGs were minor constituents, typically contributing <5-10% relative abundance per sample (Fig. 3C). As in the read-based analysis, MTZ stood out with a relatively high NRE fraction, with some nodules containing 15-25% non-*Bradyrhizobium* taxa, especially *Novosphingobium* and *Burkholderia*. Consistent with this, alpha-diversity analyses of both assembled and unassembled datasets identified MTZ as the most diverse treatment (Fig. S2C), whereas RH was the least diverse and contained only *Bradyrhizobium* MAGs. Notably, despite these differences in diversity, both RH and MTZ showed the highest fixation rates, leaf nitrogen concentrations, and plant biomass (Fig. 1B), suggesting that NRE abundance and diversity alone are not a major determinants of plant performance.

**Figure 3:**
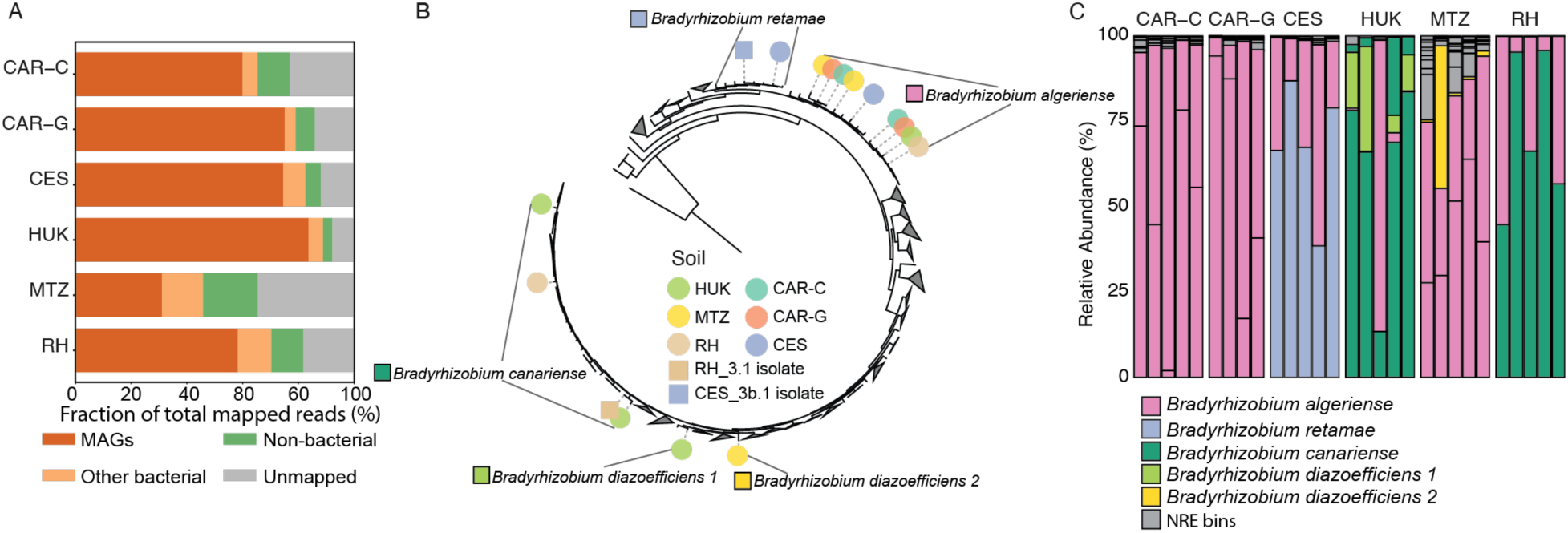
Assembly-based taxonomic Composition of Nodule Microbiomes by Site. **(A)** read mapping distribution to MAGs, non-MAG bacteria, unbinned contigs classified as non-bacterial (non-bacterial) and unmapped. **(B)** Phylogenetic tree based on bacterial house-keeping genes, of the entire *Bradyrhizobium* genus (509 genomes), including the symbionts assembled from the nodule metagenomes. Clades that did not contain our MAGs are collapsed. Species grouping of MAGs are denoted on the tree. **(C)** Bar plots showing the relative abundance of the bins found in each soil treatment based on CoverM relative abundances.

To classify the *Bradyrhizobium* MAGs, we placed them within a single-copy-gene phylogeny tree of the broader *Bradyrhizobium* genus (Fig. 3B). This analysis indicated that the symbionts in our study fall into five putative species: *B. algirense, B. canariense, B. retamae* and two distinct *B. diazoefficiens* clades. Among these, *B. algirense*, represented by nine MAGs, emerged as the dominant symbiont of C. *villosa*, as it was detected at high abundance in all treatments. In the CAR treatments, *B. algirense* was the only symbiont species recovered (Fig. 3C). Yet despite its broad presence, it was replaced by other species in some sites: CES was dominated by *B. retamae*, whereas HUK and RH were dominated by *B. canariense*. Thus, *C. villosa* nodules are colonized by multiple distinct *Bradyrhizobium* populations whose identities are shaped by soil inoculum source.

One clade, *B. diazoefficiens* 1, appeared to be strongly mutually exclusive with the primary symbiont, *B. algirense* (Fig. 3C), raising the possibility of antagonistic interactions between these two groups. To examine this pattern more closely, we used a compendium of 1,963 global soil metagenomes that we recently curated [62] and mapped the global biogeography of *Bradyrhizobium* and its major clades. We found that the *Bradyrhizobium* is the most abundant bacterial genus in soil, globally. It is a cosmopolitan soil colonizer, detected in all 1,963 soil metagenomes, with a median relative abundance of 5.49% of classified reads (Fig. S3A). *Bradyrhizobium* abundance was most strongly positively associated with mean annual precipitation (ρ = 0.48) and most strongly negatively associated with soil pH (ρ = −0.45), indicating that it reaches its highest relative abundances in humid, acidic soils rather than in the dry, alkaline soils used in this study (Fig. S3B). At the clade level, distinct environmental niches emerged: *B. algirense* and *B. canariense* were negatively associated with temperature, suggesting a preference for cooler or more seasonal climates, whereas the *B. diazoefficiens* clades, and to a lesser extent *B. retamae*, were positively associated with mean temperature of the coldest quarter and with annual precipitation, consistent with adaptation to warmer and wetter environments (Fig. S3C). Together, these results identify precipitation and pH as the main macroecological correlates of *Bradyrhizobium* abundance, while also showing that individual clades partition temperature and moisture niche space in distinct ways.

Co-occurrence analysis using FastSpar [55] revealed a broadly positive association among most *Bradyrhizobium* species, with the notable exception of *B. diazoefficiens* 1 (Fig. S3D). At the level of individual species, *B. diazoefficiens* showed its strongest negative correlation with *B. algirense* (Fig. S3D), mirroring the pattern observed in our nodule dataset and lending further support to the idea that these two groups may antagonize one another.

### Symbiont nitrogen metabolism genes predict host nitrogen status and nodule investment

To identify the nodule bacterial genes most closely associated with plant phenotype, we first carried out a targeted analysis of nitrogen metabolism genes (Table S2) against the measured plant traits. Leaf nitrogen concentration and NMF showed the strongest associations with this gene set (Fig. 4). As expected, the core nitrogen fixation genes of the *nif* operon were found predominantly in *Bradyrhizobium* symbiont MAGs (Fig. S4, S5). However, several *Bradyrhizobium* MAGs lacked part or all of the *nif* operon, even though at least one *nif*-containing MAG was present in every treatment (Fig. S5). These *nif*-deficient MAGs were associated with reduced *nif* operon KO abundance in the corresponding treatments, suggesting possible cheating behavior (Fig. S5).

**Figure 4:**
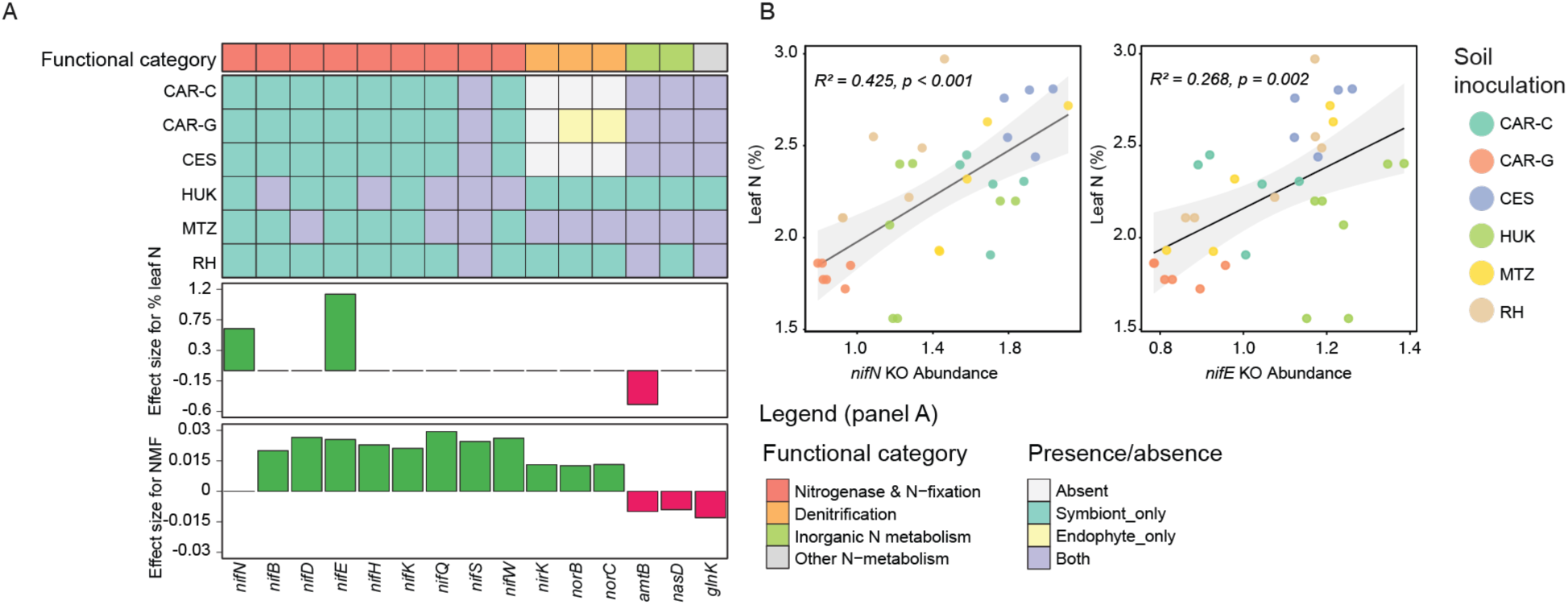
Nitrogen metabolism and fixation targeted analysis. **(A)** heatmap showing the presence of nitrogen metabolism genes that passed the targeted analysis in the metagenomically assembled bins. Bars show the effect size of each gene on the measured trait. **(B)** Scatterplot showing the correlation between plant leaf nitrogen concentration and *nifN* and *nifE* KO abundance across all treatments.

Notably, the symbiont-borne genes encoding the nitrogenase enzyme itself (*nifH*, *nifD*, *nifK*) were not the strongest predictors of leaf nitrogen concentration. Instead, the strongest signal came from accessory genes involved in biosynthesis of the iron-molybdenum cofactor (FeMo-co). In particular, *nifN* and *nifE* were the most significant predictors of leaf nitrogen concentration in the targeted nitrogen metabolism analysis (p < 0.001, R² = 0.425, and p = 0.002, R² = 0.268, respectively; Figure 4A-B), suggesting that variation in cofactor assembly capacity may be especially important for nitrogen delivery to the host. Beyond fixation itself, we found that the denitrification genes *nirK* and *norB/norC* were positively correlated with nodule mass fraction, whereas the ammonium transporter *amtB* and the P-II regulatory protein *glnK* were negatively correlated with both leaf nitrogen concentration and NMF. Together, these results suggest that nitrogen metabolism genes encoded by both the symbiont and the NREs shape plant performance in ways that extend beyond core nitrogen fixation genes.

Interestingly, while lacking *nif* genes, 70% of NRE MAGs had at least one copy of the nodulation *nodD* gene (Fig. S4, S6). The phylogeny of *nodD* genes in our experiment was generally divided into separate clades for symbionts and NREs, although one clade showed high similarity between the NREs and the main symbiont (Fig. S6). This suggests a role for *nodD* in allowing NRE coinfection with the main symbiont.

### Non-rhizobial endophyte genome content is linked to plant performance

In addition to the targeted analysis of nitrogen metabolism genes, we performed a broader search for microbial genomic predictors of plant performance using Boruta feature selection on nodule MAGs [61, 63]. This analysis identified 67 KOs as significant predictors across four plant traits: leaf nitrogen concentration, plant biomass, per-nodule biomass fixation rate, and NMF (Fig. 5). To focus interpretation, we narrowed this list to KOs that met three criteria: they belonged to relatively compact KEGG modules (<15 KOs), were significantly correlated with the relevant trait, and were present in our MAGs. Starting from these 18 selected “seed” KOs, we then examined the remaining components of their respective modules for associations with plant traits (Fig. S7). The module with the largest number of significantly correlated KOs was pyrimidine degradation, in which *rutC* was the selected seed and *rutADE* and *yfdG* were significantly and negatively correlated with NMF (Fig. S7). Another strongly associated module was assimilatory sulfate reduction, in which *cycN* was the seed gene and *cysD* and *cysNC* were negatively correlated with leaf nitrogen concentration (Fig. S7, S8). Genes in the trans-cinnamate degradation module (trans-cinnamate → acetyl-CoA) were also associated with leaf nitrogen concentration: *mhpB* was the seed gene, *mhpB* and *mhpC* were positively correlated with leaf nitrogen concentration, and *mhpA* was negatively correlated (Fig. S7). In the denitrification pathway, *norB* was selected as the seed, and although none of the denitrification genes were significantly correlated with leaf nitrogen concentration, *norBC* and *nirK* were positively correlated with NMF, consistent with the targeted analysis (Fig. 4, S7). Notably, many of the selected genes were found primarily in NREs (Fig. 5, S8). Together, these results suggest that genomic functions encoded by NREs are informative predictors of plant performance despite their low relative abundance within nodules.

**Figure 5.**
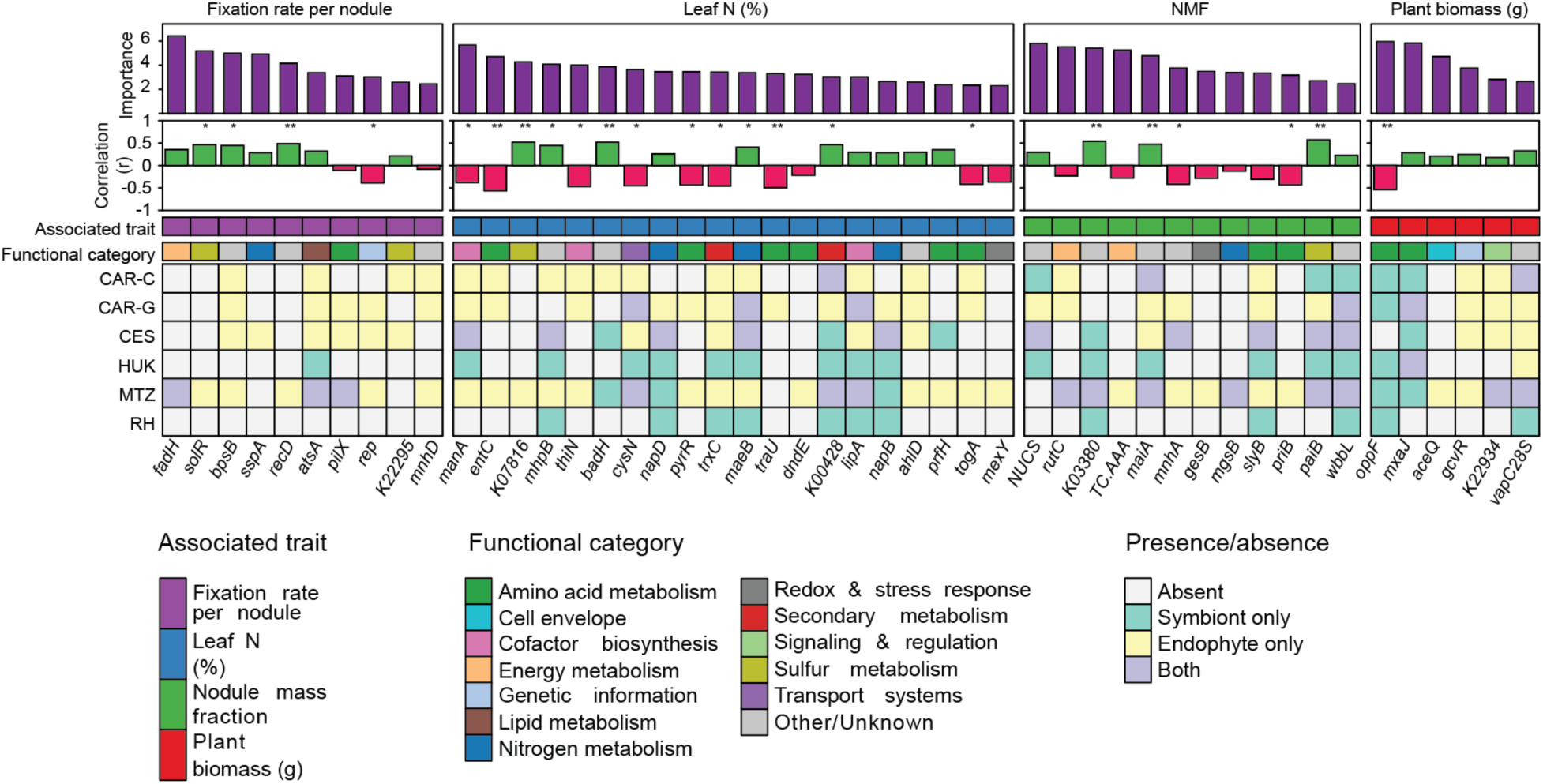
Unrestricted selection analysis of microbial genes associated with plant traits. Boruta-selected KEGG orthologs (KOs) associated with plant performance traits and their distribution across symbiont and NRE MAGs. Heatmap displaying the 67 KOs selected by Boruta analysis as significant predictors of plant performance traits across the six inoculation treatments. The main heatmap body shows for each KO whether it was detected in symbiont MAG only, NREs only, both niches, or absent from bins in that treatment. KOs are split into four column groups by the plant trait for which they were Boruta-selected: plant biomass (g), leaf nitrogen concentration (%), nodule mass fraction (NMF), and fixation rate per nodule biomass. Within each trait group, columns are ordered first by Boruta mean importance score (descending) and then by functional category. Column labels show the KEGG functional definition of each KO (truncated to 50 characters). Five annotation tracks above the main heatmap are: (i) Boruta Importance: a barplot of the mean Boruta importance score assigned to each KO across all traits for which it was selected. (ii) Correlation (r): a barplot of the Pearson correlation between per-sample KO abundance and the Boruta-associated plant trait; bars are green for positive correlations and red for negative correlations; asterisks above bars denote statistical significance (* p < 0.05, ** p < 0.01, *** p < 0.001). (iv) Associated Trait: color strip indicating which plant trait each KO was selected for plant biomass, leaf N%, nodule mass fraction and fixation rate per nodule biomass. (v) Functional category: color strip showing the KEGG-derived functional category of each KO, assigned by matching KEGG pathway descriptions and BRITE hierarchy terms to 13 predefined functional groups.

### A *Bradyrhizobium* Type VI secretion system is negatively correlated with NRE prevalence

To test whether any *Bradyrhizobium* genes were associated with NRE prevalence and diversity, we applied Boruta feature selection to sample-level alpha-diversity metrics. This analysis identified a significant association between components of the type VI secretion system (T6SS) in *Bradyrhizobium* MAGs and both nodule microbiome alpha diversity and relative NRE abundance (Fig. S9A-B). Among these components, the T6SS accessory gene *vasJ* (K11910) ranked among the top predictors across all three diversity metrics and was consistently negatively correlated with both microbiome diversity (Fig. S9A) and NRE abundance (Fig. S9B). These T6SS genes were only present in a subset of symbiont bins, mainly those classified as *B. algeriense* (Fig. S9C). Genomic reconstruction of the T6SS locus in each of these six *vasJ*-carrying bins confirmed the presence of a complete and structurally conserved secretion apparatus (Fig. S9C). Together, these results raise the possibility that symbiont-borne interference mechanisms help maintain nodule purity, although whether this ultimately benefits or harms plant performance remains unresolved.

## Discussion

Our results show that plant performance cannot be fully explained by rhizobial symbionts and their canonical symbiosis and nitrogen fixation genes alone, but instead reflects the collective functional capacity of the nodule microbiome. Although the N_2_-fixing symbiont *Bradyrhizobium* remains the dominant occupant of *C. villosa* nodules, variation in symbiotic outcomes was better captured by the broader functional composition of the microbial community.

Within *Bradyrhizobium*, species-level turnover was high both within and between inoculation treatments. Some strains from the same site were placed far apart on the *Bradyrhizobium* phylogeny and, in some cases, clustered more closely with geographically distant strains than with local ones. Our global soil metagenome analysis further showed that precipitation, including both mean values and seasonality, together with soil pH, define the main axes of niche separation among *Bradyrhizobium* clades. *Bradyrhizobium algeriense* and *B. canariense* were associated with cooler, more stable environments, with *B. algeriense* also linked to drier conditions. By contrast, the *B. diazoefficiens* clades, and the *Bradyrhizobium* genus more broadly, were more abundant in warm, seasonal, and relatively acidic soils. These patterns suggest that *B. algeriense* is better adapted to the local soil and climate conditions of our Mediterranean ecosystem, whereas *B. diazoefficiens* may be near the edge of its geographical range. The clade we identified as *B. diazoefficiens* shows anomalous behavior both in our nodule data and in the global soil dataset, compared with the genus as a whole, which appears to be driven by a negative biotic interaction with *B. algirense*. The mechanistic basis of this dynamic and its ecological implications remain to be resolved.

The nodule microbiomes extend beyond the symbionts themselves, ranging from nearly pure symbiont communities to nodules containing substantial fractions of NREs. The presence of *nodD* homologs in many NRE genomes is intriguing in this context. The *nodD* gene encodes a flavonoid-activated transcriptional regulator that induces expression of nod genes such as *nodABC*, functioning primarily as a contact-dependent activator of biosynthesis of nod factors, that in turn initiate root infection and nodule formation on compatible host plants [6, 64, 65]. In our dataset, the *nodD* phylogeny contained symbiont-only, NRE-only, and mixed clades, suggesting a more complicated evolutionary and ecological history than a strict separation between symbiotic and non-symbiotic taxa. However, whether *nodD* contributes directly to NRE entry into nodules remains unresolved.

One mechanism that may shape nodule community composition is bacterial interference by the symbiont itself. The negative correlation between components of the T6SS, and within nodule alpha diversity and NRE relative abundances suggests that symbionts can actively regulate the composition of the nodule microbiome. First discovered as a critical component used by *Rhizobium leguminosarum Biovar trifolii* to infect pea plants [66, 67], T6SS enables diverse strains to deploy toxic effectors against rival microbiota, thereby securing a competitive advantage for root nodule occupancy [68, 69] and protection from co-infection [70, 71]. Notably, T6SS was shown to play a similar role in the analogous *Vibrio fischeri-Euprymna scolopes* (bobtail squid) symbiosis [72]. At the same time, T6SS effects are host dependent: in some rhizobia it impairs effective nodulation, whereas in others it enhances nodulation success and host performance [69, 73, 74]. In our system, the negative association between T6SS abundance and nodule diversity suggests that this machinery may help maintain symbiont-dominated nodules.

Although most *nif* gene abundances per genome were positively correlated with plant NMF, only *nifN* and *nifE* were directly correlated with leaf nitrogen concentration. These two genes encode the FeMo cofactor cluster, a critical and rate-limiting step in the assembly of the nitrogenase enzyme, suggesting this is a major bottleneck for nitrogenase biosynthesis and, by extension, the supply of fixed nitrogen to the plant [75, 76]. Looking beyond fixation, our data suggests that nitrogen loss also plays a role in nodule microbiota. The denitrification pathway genes encoding nitrite reductase (*nirK*) and nitric oxide reductase (*norBC)* were positively associated with NMF. One possible explanation for this is that legume nodules are prone to the accumulation of the radical nitric oxide (NO), that can irreversibly inhibit nitrogenase and trigger premature nodule senescence, and bacterial *norBC* is the primary mechanism for removing excess NO [77]. Another explanation is that since nitrate suppresses nodulation via the autoregulation of nodulation (AON) pathway, bacterial denitrification could attenuate this suppression by removing local nitrate, inducing greater nodule investment under identical external nitrogen supply [78–80]. We also found that the ammonium transporter (*amtB*) and the P-II regulatory protein (*glnK*) are negatively correlated with both leaf nitrogen and NMF. In free-living bacteria, *amtB* is typically upregulated under nitrogen limitation to assimilate ammonium for cellular biosynthesis [81]. However, in a functional nitrogen-fixing bacteroid, *amtB* activity should be repressed to favor the export of ammonium to the host [82]. The negative association between *amtB* abundance and host nitrogen status therefore suggests that bacterial nitrogen retention may come at the expense of host benefit. Furthermore, *amtB* is present across both primary symbiotic *Bradyrhizobium* and non-rhizobial endophyte MAGs, demonstrating that both the primary symbiont and NREs, can potentially compete with the plant for symbiotically-fixed ammonium. This interpretation is also consistent with previous work showing that *amtB*-type transporters are downregulated during bacteroid differentiation, as the bacterial lifestyle shifts away from ammonium scavenging and toward nitrogen release to the plant [83–87]. Together, our findings suggest that genes that cycle nitrogen within nodules, beyond fixation, are significantly correlated with host phenotype.

An additional module related to nitrogen scavenging, the pyrimidine degradation module (*rutABCDEFG*), emerged from the untargeted analysis. This system, which enables growth on uracil or uridine as the sole nitrogen source [88, 89], was negatively correlated with NMF. One possible explanation is that local ammonium release from pyrimidine degradation contributes to nitrogen sufficiency signals that dampen nodule investment through AON. The untargeted analysis also identified a sulfur assimilation module that was negatively correlated with leaf nitrogen. Sulfur is a critical component of both nitrogenase and nodular ferredoxin, making it a potentially limiting element in the legume-rhizobia symbiosis [90]. Yet in our dataset, sulfur assimilation genes were negatively associated with leaf nitrogen concentration and were found mainly in NREs, particularly in the low-performing CAR-G treatment. A plausible interpretation is that these NREs compete with the symbiont for available sulfur, thereby constraining nitrogen fixation efficiency and host nitrogen gain [90–92]. Although this requires further validation, it illustrates how low-abundance community members may influence host function by competing for key metabolites required for efficient symbiosis.

Taken together, our data suggest that nodules are multipartite microbial communities in which cooperation, competition, and potential conflict coexist. Wild legumes growing in unmanaged soils may retain microbial associations and functional diversity that have been reduced by crop domestication and agricultural management, making them valuable systems for discovery, which may help build the knowledge needed for climate-smart agriculture that makes optimal use of biological nitrogen fixation.

## Data availability

All scripts used to do analyses and create the figures are in https://github.com/Naturalist1986/Metagenomic_Assembly_and_Binning_pipeline and https://github.com/Naturalist1986/nodule-microbiome-analysis-scripts. All data generated during the study are included as supplementary information files. Sequencing data and MAGs have been deposited in the NCBI Sequence Read Archive (SRA) under BioProject PRJNA1438069.

## Acknowledgments

We thank Prof. Yuval Gotlieb for help with nodule DNA extraction and Prof. Gabriel Castrillo for providing comments on the manuscript. The study was funded by Israel Science Foundation (ISF) grants #508/16 and #1142/19 to ES; ISF grant #2039/21, European Research Council (ERC) grant #101077278 and Binational Science Foundation (BSF) and US National Science Foundation (NSF) grant under awards #2024617 and IOS #2421771 to OMF.

## Supplementary figures

**Supplementary Figure S1:**
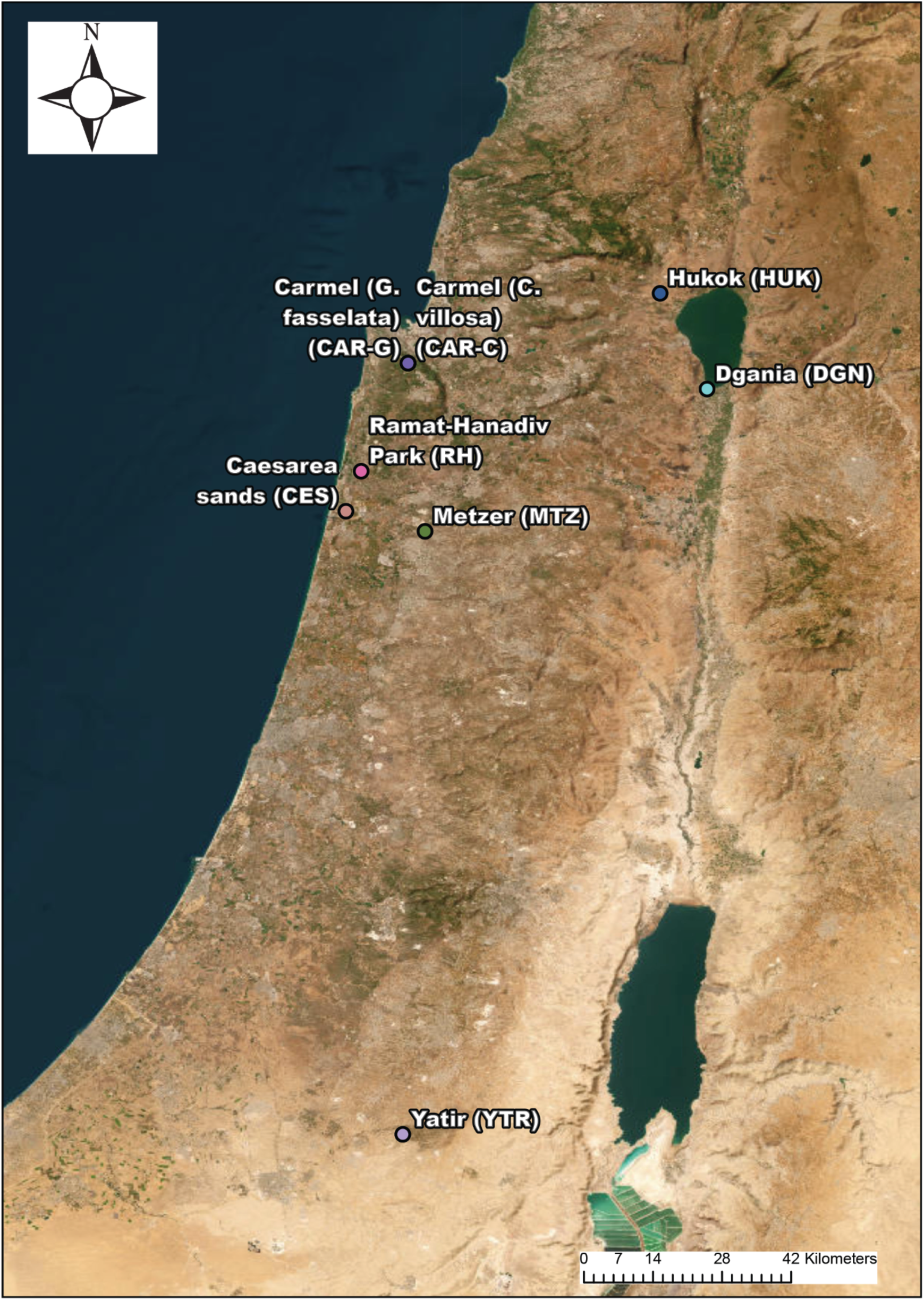
Geographical distribution of the field sites from which the soil inoculants were collected.

**Supplementary Figure S2:**
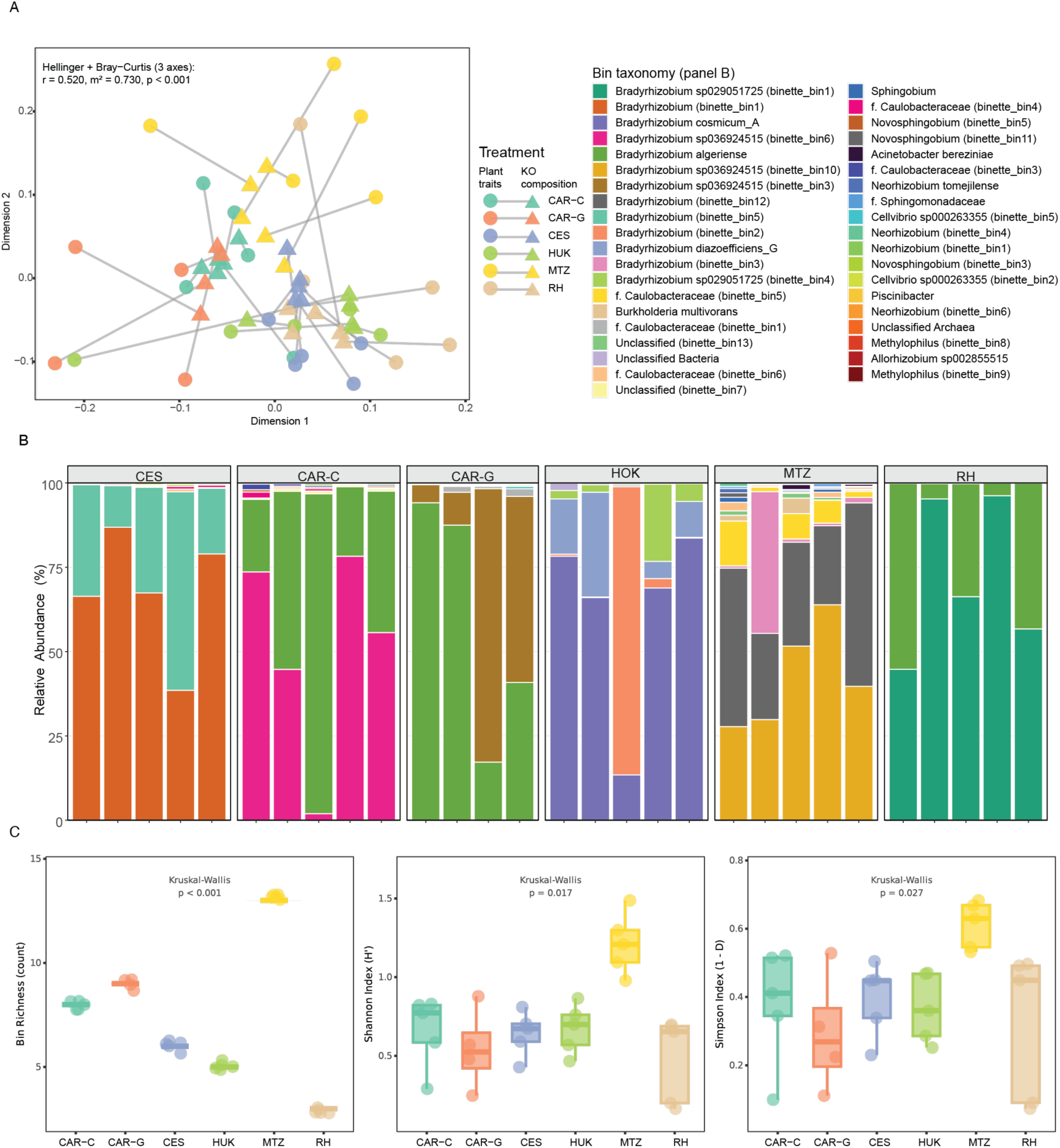
Procrustes analysis ordination and diversity measures of nodule microbiomes. **(A)** Procrustes superimposition of plant traits and KO functional composition ordinations. Filled circles represent plant trait PCA scores, filled triangles represent rotated KO composition PCoA scores (Hellinger-transformed, Bray-Curtis dissimilarity, 3 axes), colored by soil inoculation treatment. Gray lines connect matched samples, with line length indicating Procrustes residual (fit quality). Shorter lines indicate better correspondence between trait and functional composition. Analysis used a filtered dataset (7,047 KOs) with Hellinger transformation and Bray-Curtis dissimilarity (r = 0.520, p < 0.001). **(B)** Relative abundance barplot of metagenomically assembled gnomes (MAGs) based on CoverM abundance mapping. Bars show only the mapped fraction of the metagenomic reads, with abundance renormalized to add up to 100. **(C)** Diversity measures of nodule microbiomes based on MAG relative abundances. P-values for Kruskal-Wallis test by ranks are shown within each plot.

**Supplementary Figure S3:**
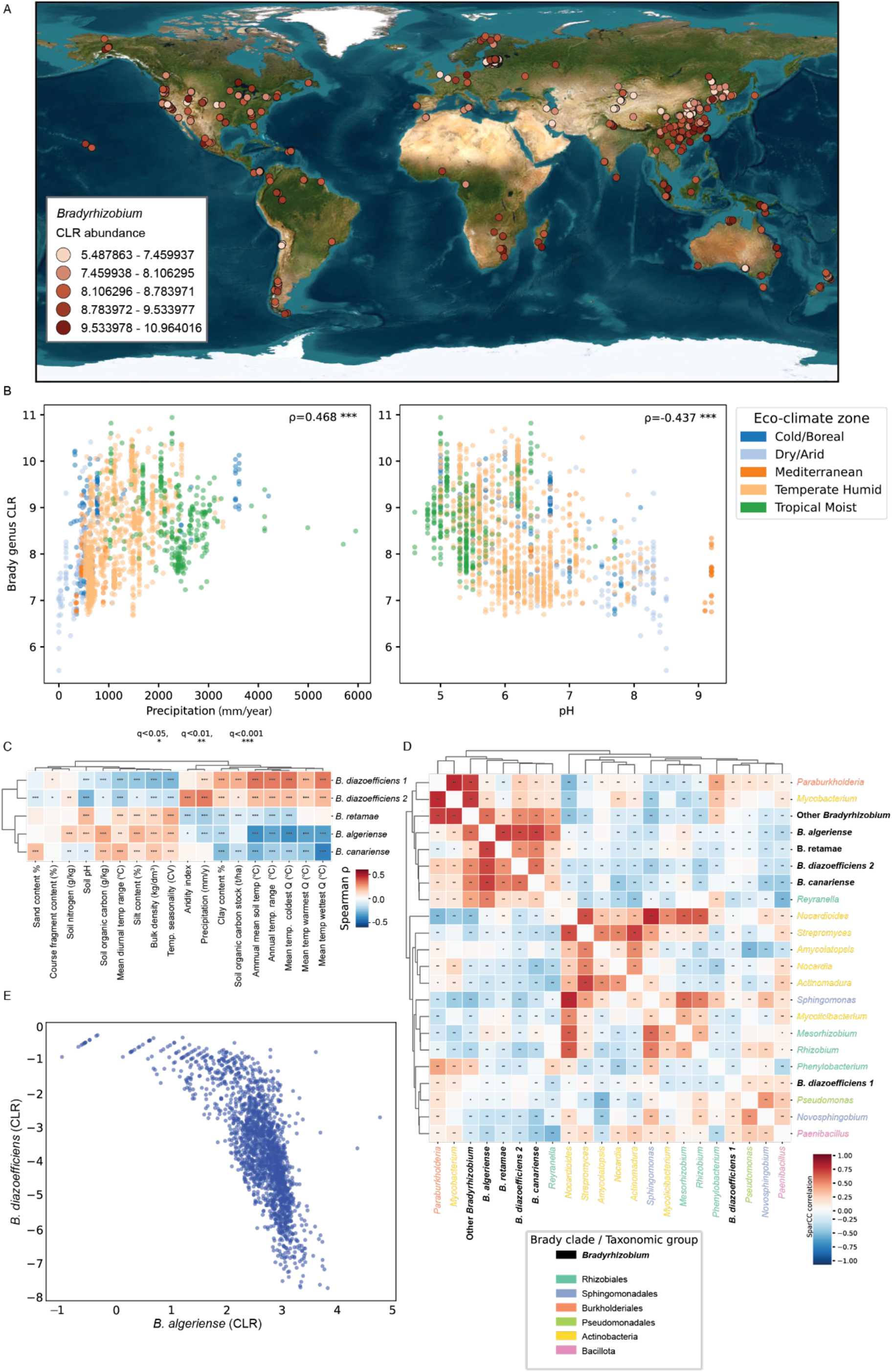
Co-occurrence of the *B. algeriense* and *B. diazoefficiens 1* clades in global soil metagenomes dataset. **(A)** CLR abundance of the *Bradyrhizobium* genus in the global soil metagenomes dataset. **(B)** CLR abundance of the *Bradyrhizobium* genus vs mean annual precipitation and soil pH. **(C)** Correlation heatmap between CLR abundance of the *Bradyrhizobium* clades and soil properties and climatic variables (ArcGIS Living Atlas, [93–95]). **(D)** Fastspar [55] co-occurrence heatmap of the *Bradyrhizobium* clades as defined in Fig. 3 and all non-bradyrhizbium genus that had ≥0.01% mean relative abundance across all samples, combined with a prevalence filter of present in ≥10% of samples **(E)** CLR-transformed clade abundance values from the 1,742 soil metagenomes of *B. diazoefficiens 1* vs. *B. algerianese.* In all panels, * q < 0.05, ** q < 0.01, *** q < 0.001.

**Supplementary Figure S4:**
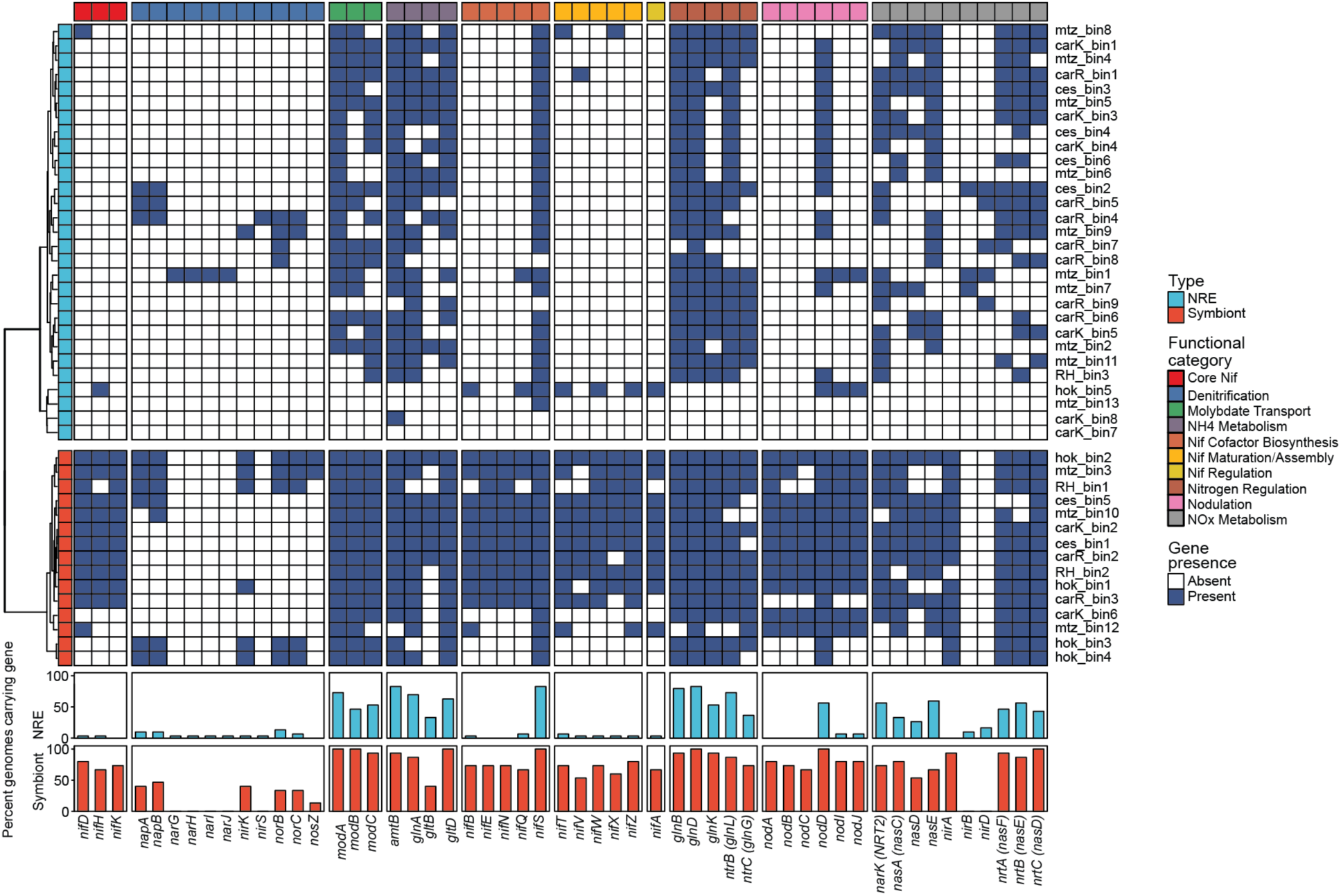
Presence/absence of nitrogen metabolism genes in MAGs. The top heatmap shows endophyte MAGs while the bottom one shows symbiont MAGs. Barplots show the percent of genomes of each MAG type (symbiont or endophyte) carrying the specific gene.

**Supplementary Figure S5:**
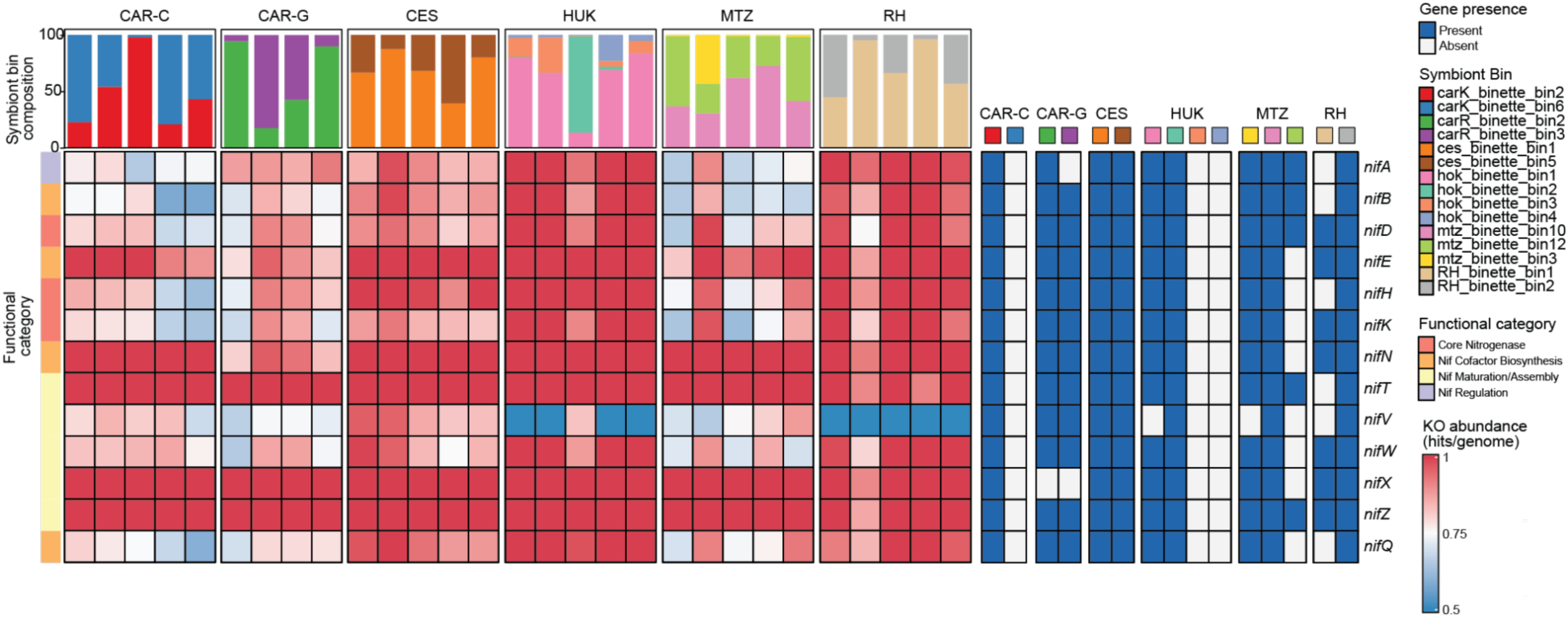
Heatmap of *nif* operon gene abundance based on KO assembly free mapping. Top barplot shows the relative abundances of the symbiont bins within the treatment according to CoverM mapping. The heatmap on the right shows presence/absence of the *nif* operon genes in the symbiont bins.

**Supplementary Figure S6:**
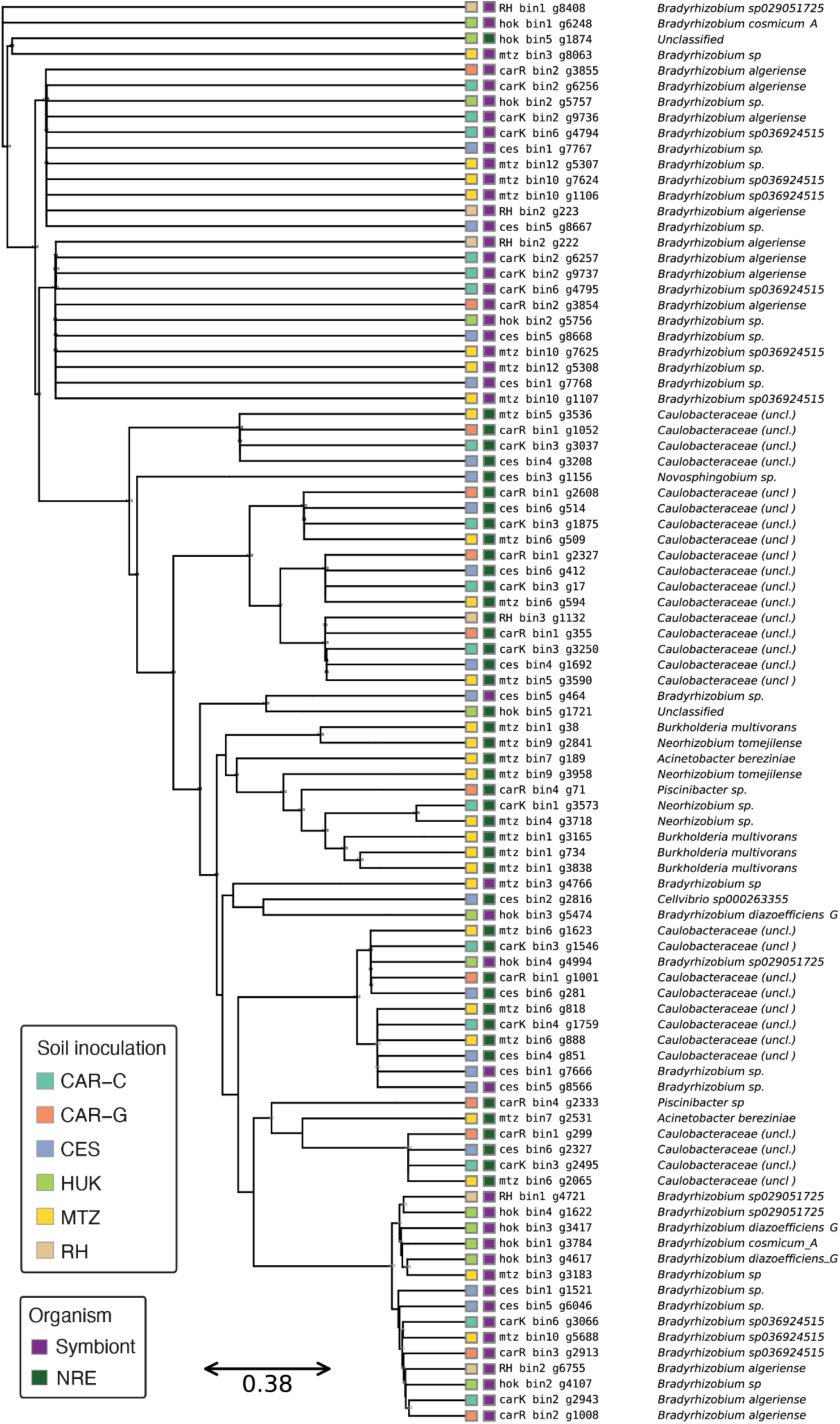
Phylogenetic tree of the nodD gene in all MAGs carrying it. MAGs may appear more than once when they carry multiple nodD gene copies.

**Supplementary Figure S7:**
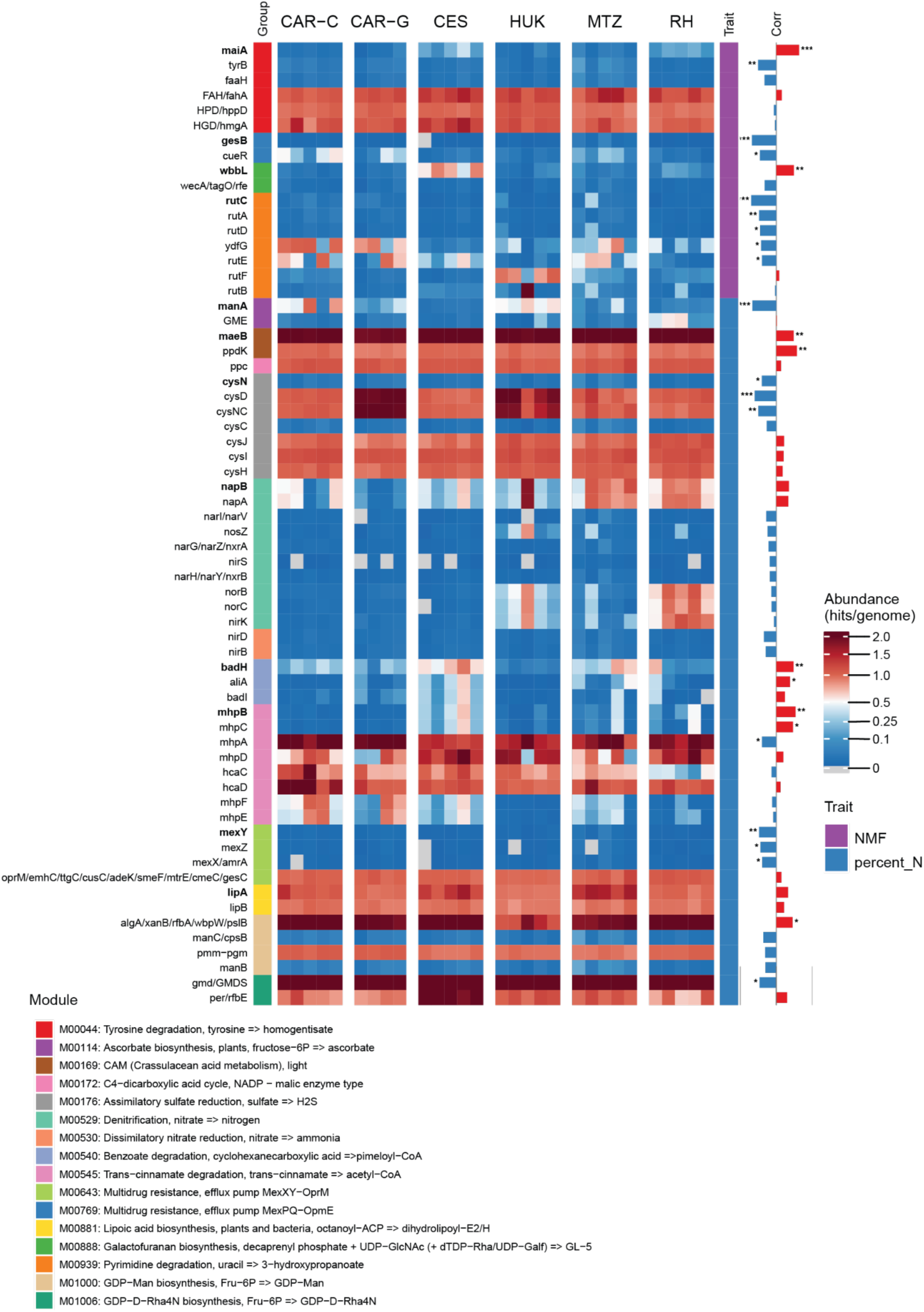
KEGG module context of Boruta-selected KOs: co-occurring pathway members and their associations with plant performance traits. Heatmap displaying per-sample normalized abundances (hits per genome) for all KEGG orthologs (KOs) belonging to KEGG modules that contain at least one Boruta-selected KO. Boruta KOs served as seeds: for each seed, all KOs in the same module that were detected in at least one sample were included as companion KOs. Only modules with ≤10 KOs detected in the dataset and at least one non-Boruta companion KO were retained, excluding large overview modules. Rows are not clustered but are arranged hierarchically: first by the plant trait associated with the seed KO (leaf N concentration, plant biomass, fixation rate per nodule biomass, nodule mass fraction), then by module, and within each module seeds appear before companions, with companions ordered by the absolute magnitude of their Spearman correlation with the seeded trait. Bold labels indicate Boruta seed KOs. Corr: a horizontal barplot showing the Spearman correlation coefficient (r) of each KO with the seeded trait, with bars extending right (red) for positive correlations and left (blue) for negative correlations; significance asterisks mark bars with p < 0.05 (*), p < 0.01 (**), and p < 0.001 (***).

**Supplementary Figure S8:**
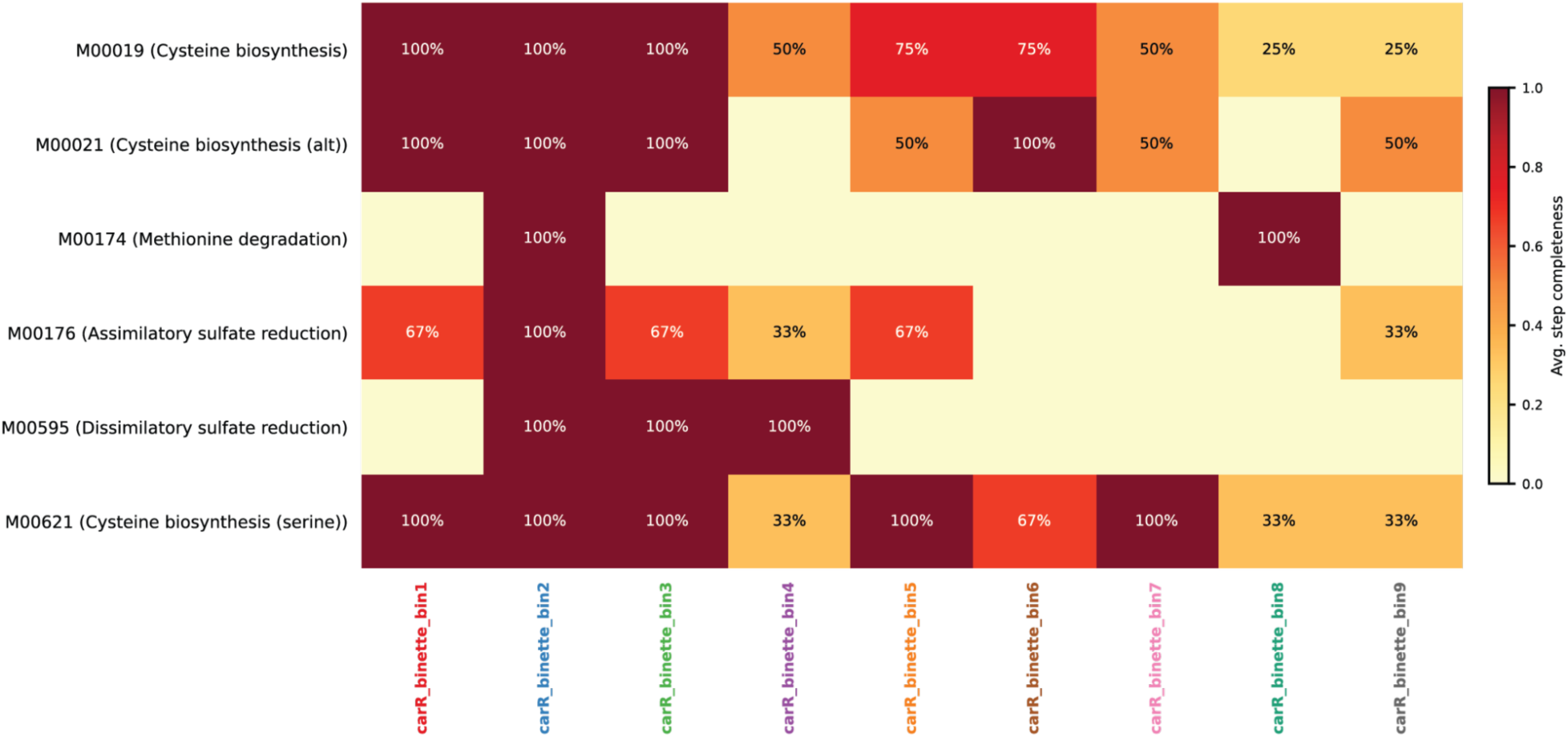
Sulfur metabolism in CAR-G treatment. Pathway completeness of sulfur metabolism modules within each of the MAGs obtained from the CAR-G treatment.

**Supplementary Figure S9:**
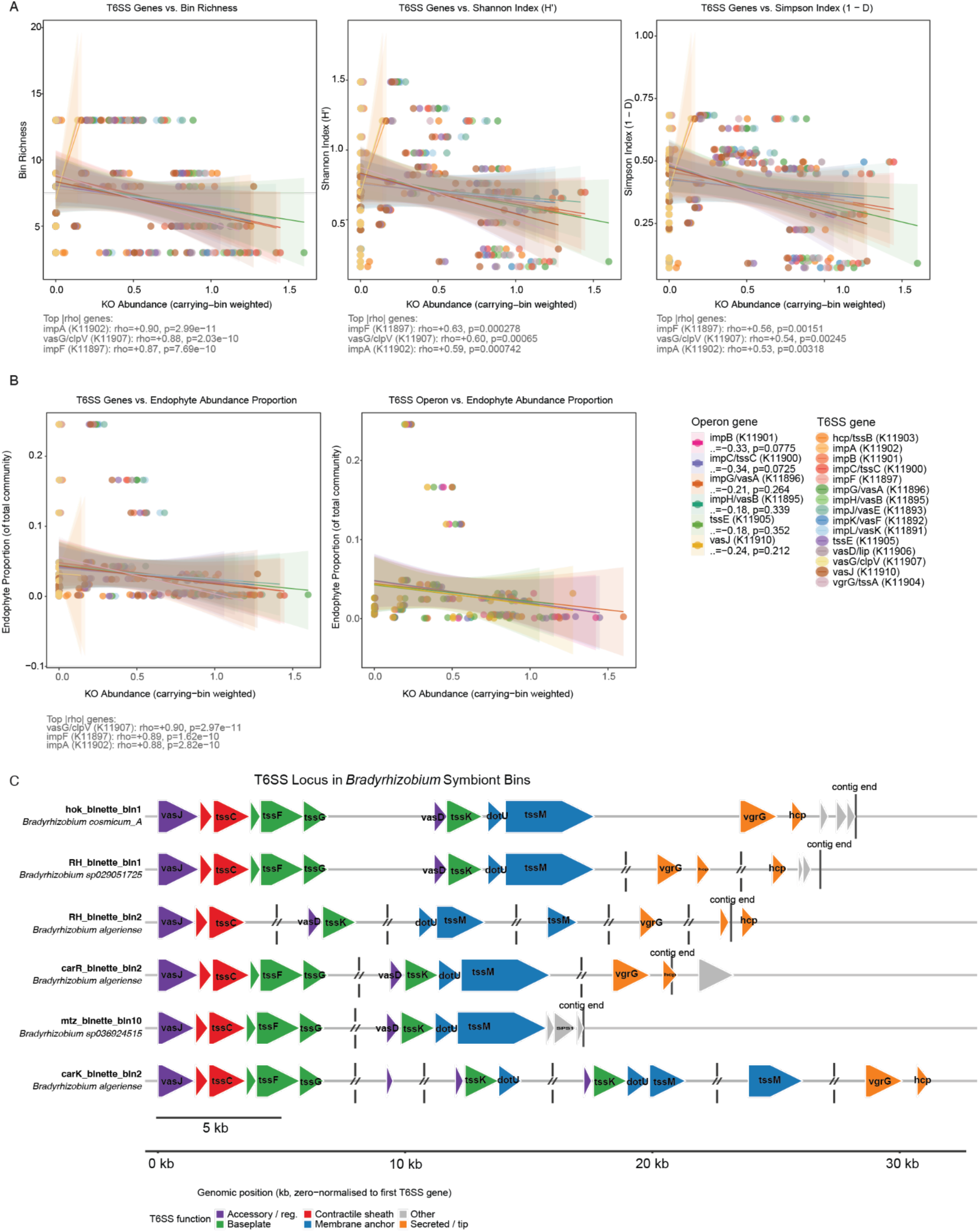
Type VI secretion system locus in Bradyrhizobium symbionts. **(A)** Regressions of alpha diversity measures against per genome abundances of the different components of the type VI secretion system. **(B)** Regressions of NRE relative abundance against per genome abundances of the different components of the type VI secretion system. **(C)** Structure and components of the type VI secretion system in the six symbiont MAGs that carry it. Vertical line and slashes show contig breaks, likely due to a fragmented assembly. Each bin harbours genes encoding all four functional modules: the contractile sheath (*impB/tssC*), the baseplate complex (*tssE, tssF, tssG, tssK*), the membrane anchor (*dotU, tssM*), and the accessory-regulatory components (*vasJ, vasD*), as well as the secreted effector-delivery proteins (*hcp* and *vgrG*) (Fig. S9C). The complete locus spans approximately 8-26 kb of genomic sequence and, in four of the six bins, was distributed across two to three contigs as a result of assembly fragmentation at the *tssG/vasD* junction. Despite this fragmentation, the canonical gene order was preserved in all bins. The bin with the most complete contiguous assembly (*B. cosmicum A*, HUK treatment) encoded all 12 core T6SS KOs on a single scaffold spanning 25.8 kb, confirming operon integrity.

## Notes

https://github.com/Naturalist1986/Metagenomic_Assembly_and_Binning_pipeline

https://github.com/Naturalist1986/nodule-microbiome-analysis-scripts

https://www.ncbi.nlm.nih.gov/bioproject/?term=PRJNA1438069

### Competing Interest Statement

The authors have declared no competing interest.

