## Supplementary Methods for "Nodule microbiome functions shape the performance of a wild perennial legume"

### Metagenomic Analysis

Metagenomic sequencing data from nodule samples were processed using a metagenomic assembly and binning pipeline available at github. Raw paired-end reads were quality filtered using Trimmomatic with adapter trimming and a minimum read length of 36 bp, followed by read validation and repair using BBTools [1, 2]. From this point, to gain a comprehensive image of nodule community composition, we combined assembly-free and assembly-based annotation approaches. In the assembly-free approach, taxonomic and functional profiles are obtained directly from unassembled sequence reads. This provides an image of the community composition that is not biased by the variable success of assembly. In the assembly-based approach, reads are assembled and binned into metagenomically assembled genomes (MAGs), and abundances are calculated by mapping reads back to these assemblies. This enables an analysis that considers genomic context, and resolving the genomes of symbionts and NREs, as well as distinguishing closely related strains within the genomes. MAGs are available at bioproject PRJNA1438069 at NCBI.

### Assembly-free Functional Annotation and KEGG Mapping of Nodule Metagenomes

We ran an assembly-free analysis of the nodule metagenomes. Metagenomic reads from nodule samples were quality filtered using Trimmomatic v0.39-2 with adapter trimming (ILLUMINACLIP with 2:30:10 parameters) and quality-based trimming (SLIDINGWINDOW:4:20, MINLEN:50). Taxonomic profiles were generated using SingleM with the GTDB r226 reference database [3]. Quality-filtered reads were functionally annotated by mapping to the KEGG Prokaryotes database using DIAMOND BLASTX in sensitive mode with an e-value cutoff of 1e-5 and reporting the top hit per query (--max-target-seqs 1) [4]. Raw KEGG ortholog (KO) counts were normalized by estimated genome number to account for variable microbial biomass across samples. Genome counts were estimated by calculating the mean coverage of 106 bacterial single-copy KEGG genes per sample [5]. The resulting genome-normalized feature table was used for downstream functional analysis and for comparative metagenomics.

### Assembly, Bin Recovery and MAG annotation

Quality-filtered reads were assembled using MetaSPAdes with default parameters optimized for metagenomic data [6]. Assembled contigs were binned using five binning algorithms: MetaBAT2 , MaxBin2 , CONCOCT, COMEBin and SemiBin2 [7–10]. Bins from all five methods were integrated and refined using Binette to generate consensus bin sets [11]. Bin quality (completeness and contamination) was assessed with CheckM2 [12]. The highest quality version of each bin was automatically selected based on a quality score calculated as completeness minus five times contamination. Relative abundance of each metagenomically-assembled genome (MAG) across all samples was quantified using CoverM with the trimmed mean method [13]. Taxonomic classification of MAGs was performed using GTDB-Tk against the Genome Taxonomy Database (release 226) . Only high-quality (≥90% completeness, ≤5% contamination) and medium-quality (≥50% completeness, ≤10% contamination) bins, following

MIMAG standards, were retained for downstream analyses [14]. MAGs were annotated using Prodigal and Anvi'o [15, 16]. We used Anvi'o to construct a phylogenetic tree of the *Bradyrhizobium* genus including our classified *Bradyrhizobium* MAGs [16].

Metagenomic bins were screened for eukaryotic content using EukFinder [17], a two-stage classifier that combines Centrifuge-based taxonomic classification with PLAST protein homology searches against a UniProt reference database. EukFinder was then run on bins from each treatment, classifying assembled contigs into six categories: bacterial (Bact), archaeal (Arch), eukaryotic (Euk), unknown (Unk), eukaryotic-unknown (EUnk), and miscellaneous (Misc). Per-bin classification counts and sequence sizes were aggregated into a consolidated summary table spanning all treatments and binning tools. Non-prokaryotic sequences (Euk, Unk, EUnk, and Misc) were extracted from all bins and pooled per treatment. Treatment-level reads were then mapped back to these non-prokaryotic sequence pools using Bowtie2, and mapping rates were summarised to quantify the proportion of metagenomic reads attributable to non-prokaryotic organisms in each sample.

**Co-occurrence analysis of nodule-associated MAGs**

To investigate co-occurrence patterns among metagenome-assembled genomes (MAGs) recovered from nodule metagenomes, we employed FastSpar, a C++ implementation of the SparCC algorithm designed for compositional microbiome data [18]. Relative abundances of 44 MAGs were derived from 29 nodule metagenome samples spanning the six soil inoculation treatments. Relative abundances were scaled by a factor of 10,000 to generate pseudo-counts and reformatted as an OTU table with samples as columns and MAGs as rows, as required by FastSpar. Compositional correlations were estimated under two complementary frameworks: a MAG-level analysis retaining all 44 individual genomes and a clade-level analysis in which *Bradyrhizobium* MAG abundances were summed within each of the five clades, yielding 34 OTUs in total (5 Bradyrhizobium clades plus 29 non-Bradyrhizobium MAGs). Statistical significance of pairwise SparCC correlations was assessed using 1,000 bootstrap resamples to generate empirical p-values. Raw p-values were corrected for multiple comparisons across all pairwise combinations using the Benjamini-Hochberg false discovery rate (FDR) procedure, and pairs with $q < 0.05$ were considered statistically significant. Results were visualised as hierarchically clustered correlation heatmaps with significance annotations ($q < 0.05$, $q < 0.01$, $q < 0.001$).

**Association of Bradyrhizobium abundance with global soil environmental variables**

To characterise the environmental drivers of *Bradyrhizobium* abundance across global soil metagenomes, we analysed Kraken2 classification reports from 1,963 soil metagenomes curated previously [19, 20]. For each sample, reads assigned to the *Bradyrhizobium* genus (NCBI taxid 374), to each of five phylogenetic clades, and to each of 15 individual *Bradyrhizobium* MAGs were extracted and normalised by the total number of classified reads. Normalised values were subsequently transformed using the centred log-ratio (CLR), with a count-aware pseudocount of 0.5 / classified_reads applied prior to log transformation to handle zero-abundance observations while preserving the compositional structure of the data. CLR-transformed abundances were linked to 18 soil physicochemical and climatic variables obtained from the ArcGIS Living Atlas, including soil pH, total nitrogen, soil organic carbon

(SOC), bulk density, clay, sand, and silt fractions, coarse fragment content, mean annual precipitation, aridity index, mean annual temperature, diurnal temperature range, temperature seasonality, annual temperature range, and mean temperatures of the wettest, warmest, and coldest quarters [21–23]. Only soil samples for which both Kraken2 reports and complete environmental metadata were available were retained (n = 1,742). Spearman rank correlations between CLR-transformed *Bradyrhizobium* genus abundance and each environmental variable were computed and ranked to identify the strongest predictors of *Bradyrhizobium* relative abundance. Clade- and MAG-level associations with environmental variables were assessed in parallel using the same approach, with results visualised as clustered heatmaps in which hierarchical clustering was applied to both clades and environmental variables to reveal co-varying axes. MAGs were included in the MAG-level analysis only if they were detected (>0 classified reads) in more than 5% of soil samples. All p-values were corrected for multiple comparisons using the Benjamini-Hochberg FDR procedure, and associations with $q < 0.05$ were considered statistically significant.

**Statistical Analysis of Bacterial Gene-Plant Trait Associations**

**Data Integration and Preprocessing.** The normalized KO abundance of bacterial genes data were integrated with plant trait measurements including leaf nitrogen concentration (%N), total plant biomass, fixation per nodule, and nodule mass fraction. Technical replicates from multiple sequencing runs were averaged to produce single abundance values per biological replicate.

**Ordination of plant traits and bacterial composition.** To assess whether variation in plant phenotypic traits co-varies with nodule microbiome functional composition, we performed Mantel tests using the Vegan package in R [24, 25]. KO abundance profiles were first filtered to retain only orthologs present in at least 10% of samples with a mean abundance exceeding 0.0001; where duplicate sequencing runs existed for the same biological sample, abundances were averaged prior to analysis. Euclidean distance matrices were computed for each of eight plant traits (leaf nitrogen concentration, plant height, nodule biomass, plant biomass, nitrogen fixation rate per nodule mass, nitrogen fixation per plant, root mass fraction, and nodule mass fraction) and a Bray-Curtis distance matrix was computed for the filtered KO composition. Mantel tests were then performed between the KO composition distance matrix and each individual trait distance matrix, yielding a Mantel r statistic and permutation-based p-value for each trait-microbiome association. To visualize these results alongside inter-trait relationships, we generated an integrated correlation-network figure using the linkET package [26].

**Data Transformation and Ordination.** Plant trait data (four variables) were standardized to mean = 0 and standard deviation = 1 to account for different measurement scales, following standard recommendations for multivariate environmental data [27]. We used Principle component analysis (PCA) ordination to analyze the standardized plant trait variables. For KO functional composition, we tested two transformation approaches reflecting different ecological assumptions: Hellinger transformation and Bray-Curtis dissimilarity; This approach is recommended for abundance-based microbiome data where double-zeros (joint absences) should not contribute to similarity [28]. Hellinger-transformed KO abundances were subjected to Principal Coordinates Analysis (PCoA) using Bray-Curtis dissimilarities. Plant trait ordination used PCA on standardized variables. The resulting ordination configurations (plant traits vs. KO composition) were compared using Procrustes analysis.

**Evaluating correlation between the functional composition of the nodule microbiome and plant performance.** Procrustes analysis quantifies the congruence between two ordination configurations by rotating, translating, and scaling one configuration to maximize its similarity to the other [29]. We used symmetric Procrustes rotation, to match the ordination of plant traits and the nodule functional composition, which treats both matrices equally, implemented in the Vegan R package. Following Lisboa et al. (2014), we tested configurations using two, three, and four ordination axes to assess robustness of findings across different dimensionality choices (limited to four axes according to the number of plant traits) [29]. The Procrustes sum of squares ($m^2$) measures dissimilarity between configurations, with smaller values indicating greater congruence. The Procrustes correlation (r) is calculated as $r = \sqrt{(1 - m^2)}$, with values closer to 1 indicating stronger correlation. Statistical significance was assessed using PROTEST [30], which permutes rows of one matrix (9,999 permutations) to generate a null distribution of $m^2$ values under the hypothesis of no relationship between matrices. We compared four analytical approaches: (1) unfiltered KOs with Hellinger + Bray-Curtis transformations, (2) unfiltered KOs with CLR + Euclidean transformations, (3) filtered KOs with Hellinger + Bray-Curtis transformations, and (4) filtered KOs with CLR + Euclidean transformation. This comparison allowed evaluation of both filtering impact and transformation choice.

**Feature Filtering.** Following established best practices in microbiome research, we applied conservative filtering to remove rare and low-abundance KOs while retaining ecologically relevant functional genes [31, 32]. From the initial 9,031 KOs, we retained only those present in ≥10% of samples (≥3 samples) with mean relative abundance ≥0.01%. This dual-threshold approach, consistent with ANCOM-BC defaults and the Microbiome Quality Control (MBQC) project protocols, removed predominantly singleton and doubleton equivalent features while retaining 7,047 KOs (78.0% of the total) [39]. Goldmann et al. (2016) demonstrated that removing rare fungal OTUs (≤3 reads) had negligible impact on ordination-based community analyses (Procrustes correlation = 0.9964, $p<0.001$), supporting our filtering strategy.

**Software and Implementation.** All analyses were conducted in R version 4.3.3 using vegan (version 2.6-4) for ordination and Procrustes analysis, tidyverse (version 2.0.0) for data manipulation, readxl for Excel file import, and ggplot2 for visualization. Analysis scripts are available at github.

**Targeted Analysis of Nitrogen Metabolism Genes.** Given our focus on legume-rhizobia symbiosis and nitrogen fixation, we performed a targeted, hypothesis-driven analysis of genes involved in nitrogen cycling pathways. We curated a comprehensive list of 87 KEGG orthologs spanning the major branches of microbial nitrogen metabolism (Supplementary Table 2), including nitrogen fixation (core nitrogenase complex and associated cofactor biosynthesis, nitrogenase maturation, electron transfer, and regulatory genes, as well as alternative vanadium-dependent and iron-only nitrogenases) and nodulation, assimilatory and dissimilatory nitrate and nitrite reduction, dissimilatory nitrate Reduction to ammonium (DNRA), ammonia assimilation via the glutamine synthetase (GS)/glutamate synthase

(GOGAT) [33] , nitrification, denitrification, anaerobic ammonium oxidation (anammox), nitrogen transport and signaling.

For this focused gene set, we used univariate linear regression to analyze the relationship between the abundance of each KO from the list and each of the measured plant traits (leaf nitrogen concentration, plant biomass, fixation rate per nodule, and nodule mass fraction) for a total of 348 regressions. We extracted effect sizes ($\beta$ coefficients), standard errors, t-statistics, p-values, and $R^2$ values for each association. Because nitrogen metabolism genes represent a biologically coherent pathway with expected functional interdependencies, we applied four complementary correction strategies to control for false discoveries while maintaining statistical power. First, standard Benjamini-Hochberg FDR correction was applied across all nitrogen metabolism tests [34]. Second, Storey's q-value method was used to estimate the proportion of true null hypotheses ($\pi_0$), providing a measure of the minimum FDR at which each test would be called significant [35]. Third, trait-stratified FDR correction was applied independently within each trait, preventing highly significant associations in one trait from inflating significance thresholds for others. Fourth, the effective number of independent tests (M_eff) was estimated from the Spearman correlation structure of KO abundances using the Galwey method implemented in the poolr package, accounting for the non-independence of functionally related genes within nitrogen cycling pathways and yielding a less conservative correction when genes cluster into functional modules [36]. Associations were considered robust if they reached significance (adjusted $p < 0.05$) under at least two of the four correction methods. This multi-method consensus approach balances Type I error control with the discovery of true biological associations.

To understand the biological context of significant gene-trait associations, we mapped robust KOs to their taxonomic origins using the MAG data. For each nitrogen metabolism KO, we determined presence or absence in MAGs classified as either nitrogen-fixing symbionts (*Bradyrhizobium*) or NREs (other bacterial genera), and categorized each gene as symbiont-enriched, endophyte-enriched, or shared based on prevalence differences across organism types.

**Unrestricted search for functional genes for plant performance.** To identify the most predictive functional genes for plant performance, we used machine Boruta feature selection [37]. Boruta creates shadow features (permuted copies of all original features) and compares the importance scores of real features against the maximum importance among shadow features. Features that consistently perform better than the best shadow features are confirmed as truly predictive, while those that perform worse are rejected. Confirmed features represent KOs that are demonstrably better predictors than random chance, even after accounting for the complex correlation structure among genes. We annotated confirmed features with KEGG descriptions and ranked them by mean importance across iterations.

**Data Visualization**

Results were visualized using ggplot2, pheatmap, RColorBrewer, patchwork, ggrepel and associated packages [38–41]. Key visualizations included Top KO-trait correlations from genome-wide analysis (pheatmap package) [42].
